# RNA Polymerase II-mediated transcription is required for repair of ribosomal DNA breaks in nucleolar caps, guarding against genomic instability

**DOI:** 10.1101/2023.10.20.563274

**Authors:** Cintia Checa-Rodríguez, Beatriz Suárez-Quintero, Laura Contreras, Lea Korsholm, Dorthe Helena Payne-Larsen, Jesús de la Cruz, Jiri Bartek, Daniel Gómez-Cabello

**Author notes:** These authors contributed equally (Cintia Checa-Rodríguez, Beatriz Suárez-Quintero).

## Abstract

Ribosomal DNA (rDNA) double-strand breaks (DSBs) threaten genome integrity due to the repetitive and transcriptionally active nature of rDNA. The nucleolus, while central to ribosome biogenesis, also functions as stress sensor. Here, we identify a transcription-dependent mechanism in which RNA polymerase II (RNAPII) is essential for homologous recombination (HR) repair of rDNA DSBs. Using CRISPR-induced breaks, high-resolution imaging, and transcriptional inhibition, we show that RNAPII activity drives the formation of nucleolar repair caps. Mechanistically, CtIP promotes RNAPII recruitment and H3K36 trimethylation at rDNA lesions, facilitating HR. Disruption of this RNAPII–CtIP–H3K36me3 axis impairs cap formation and repair, leading to persistent damage. RNAPII inhibition exacerbates genome instability and synergizes with rDNA breaks to induce cancer cell death, without acutely impairing ribosome function. These findings uncover a co-transcriptional mechanism of rDNA repair and highlight RNAPII-mediated chromatin remodeling and spatial reorganization as key to nucleolar genome maintenance and potential targets for cancer therapy.

**Teaser:** Nascent RNA synthesis by RNAPII safeguards ribosomal DNA integrity to avoid genomic instability in human cells.

**Graphical Abstract:** 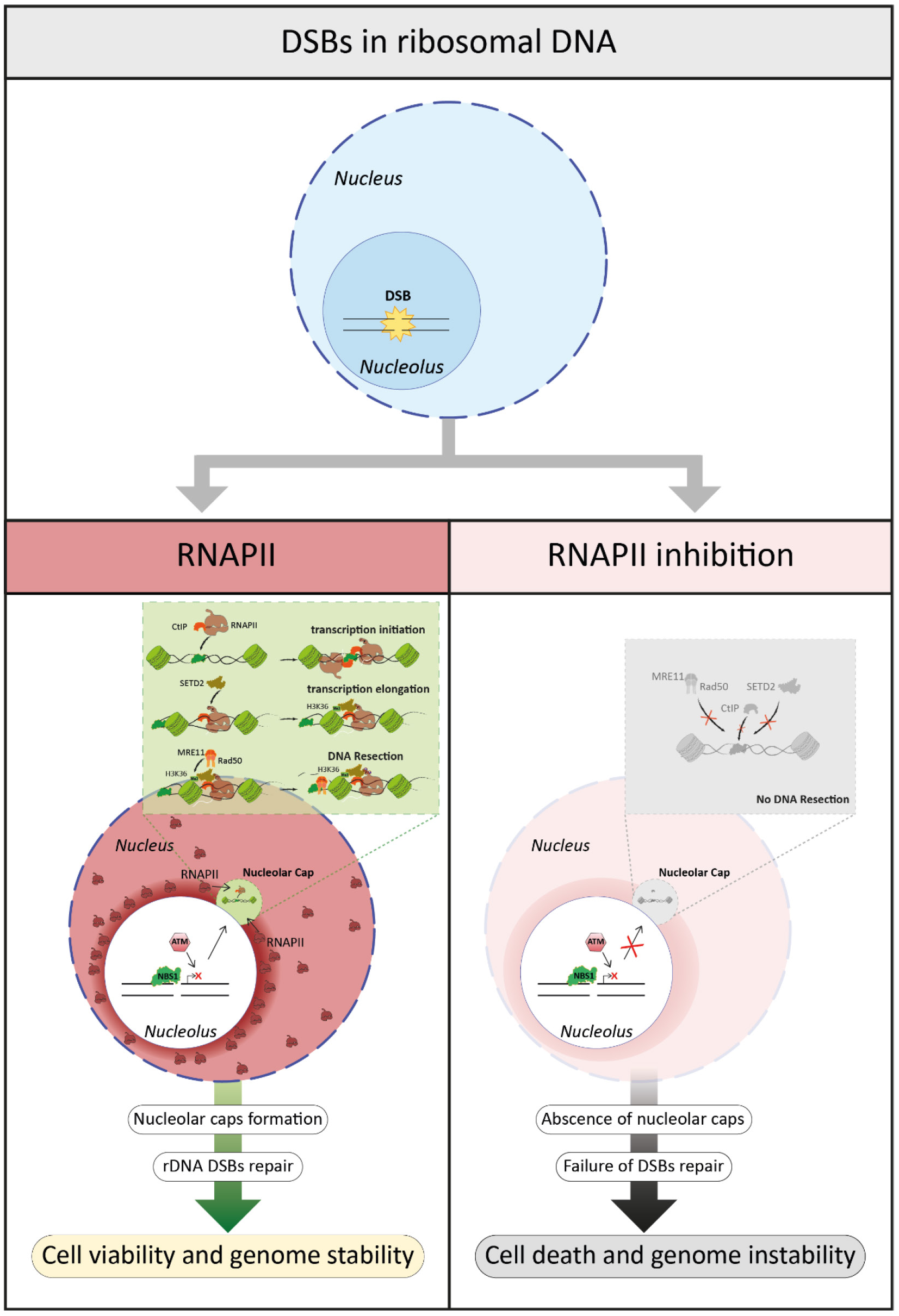

## Introduction

The nucleolus is a cellular sensor for diverse exogenous and intracellular insults. The nucleolus is a membrane-less intranuclear organelle whose canonical function is ribosome biogenesis (RiBi). RiBi includes ribosomal DNA (rDNA) transcription, rRNA processing and assembly of ribosomal subunits that are orchestrated in a complex multistep process leading to ribosome formation ^1^. Enlarged nucleoli due to elevated protein synthesis and cell metabolism, including RiBi, adapted to cope with aberrant growth and proliferation, is one of the long-established hallmarks of cancer cells ^2^. The nucleolus is also considered a central response hub for the stressors that promote cancer development ^3^. The study of nucleolar stress has been motivated, among other reasons, by efforts to interfere with RiBi function or inhibit RNA polymerase I (RNAPI), the enzyme responsible for transcription of rDNA, as strategies for cancer treatment in both preclinical and clinical settings^4^. The human genome contains around 200-600 tandem repeats of 28S, 18S and 5.8S rDNA genes dispersed in chromosomes 13, 14, 15, 21 and 22. The active rDNA sequences are concentrated in the nucleolar organizer regions (NORs), which are prone to replication-transcription conflicts potentially leading to genome instability ^5^. Such conflicts occur mainly because RNAPI-mediated transcription represents almost 60% of the total cellular transcription in eukaryotic cells ^6, 7^. The occurrence of such hazardous transcription-replication conflicts is enhanced in tumor cells which show both aberrantly elevated rDNA transcription and deregulated genome replication as common features of oncogenic transformation ^8–10^. Due to such ‘nucleolar stress’, rDNA instability occurs commonly during tumorigenesis, likely fueling tumor progression due to rDNA rearrangements events including insertions, translocations and amplifications ^11^. Among the different rDNA lesions arising from transcription-replication conflicts, the rDNA double-strand breaks (rDNA DSBs) are arguably the most cytotoxic and difficult to repair type of DNA damage ^12–14^. The two main pathways for DSB repair are homologous recombination (HR) and non-homologous end joining (NHEJ), both of which contribute to rDNA stability upon oncogenic or genotoxic stresses, such as activated cMYC oncogene and ionizing radiation (IR), respectively ^14, 15^. The NHEJ process involves little if any DNA end processing and is potentially mutagenic. On the other hand, the HR pathway is a more precise DSB repair mechanism that avoids loss of genetic information, due to faithful repair based on using a homologous DNA sequence as a template ^13, 16^. However, the HR machinery is largely absent from the nucleolar inside since intra-nucleolar recombination processes would induce illegitimate rearrangements among the many rDNA repeats localized on different chromosomes, with catastrophic consequences for the cell ^17^.

Key elements of the complex spatio-temporal cascade of DNA damage signaling and repair factors in response to rDNA lesions remain poorly understood. Recently, it has been shown that upon rDNA DSB formation, ATM-TCOF interplay promotes transient transcriptional inhibition of RNAPI and rDNA relocalization into newly formed structures at the nucleolar periphery, known as nucleolar caps ^18–20^. However, the roles of various HR factors in rDNA repair appear to be controversial, depending on the type of DSB-inducing insult, for example, IR *versus* the enzymatically induced DNA breaks ^21^. Indeed, HR factors such as RAD51 contribute to rDNA repair but their accumulation in the absence of the BLM protein, increases genome instability^22^ . Moreover, it has not yet been established what determines relocation of some rDNA DSBs to the nucleolar periphery for their HR-mediated repair, in so-called Nucleolar caps.

Recently, we and others have demonstrated the essential role of RNA polymerase II (RNAPII) and RNA polymerase III (RNAPIII) in the cell’s choice of DSB repair by HR rather than NHEJ in the whole genome. This decision is facilitated by transcription of nascent RNA flanking the DSB sites, as an early and pivotal^12, 23, 24^. Although the crosstalk between transcription and DSB repair is well established, it remains controversial whether RNAPII^25–27^ or whether its presence is limited to transcriptionally active regions, where RNA:DNA hybrids form as a consequence of ongoing transcription^28–30^. Furthermore, generally RNAPII inhibition impairs recruitment of HR factors to DSBs, mainly by preventing the early steps of CtIP^12^.

Notably, the molecular signals and mechanisms driving the nucleolar cap formation and HR engagement in repair of rDNA DSBs remain poorly defined. Until recently RNAPII was not thought to play a role in the nucleolus but it has been shown to be an important regulator of RNAPI transcription^31, 32^. More recently genotoxic stress was shown to induce RNAPII dependent transcription of the intergenic spacer region in rDNA sequestering aberrant transcripts to promote DNA repair^33^. However, if RNAPII plays a role in regulation of the specialized nucleolar DNA Damage Response (nDDR) remains unexplored. This raises the possibility that RNAPII may be locally co-opted to support rDNA repair, particularly through its transcription-coupled chromatin remodeling functions.

Here, we demonstrate that RNAPII activity is essential for the accurate repair of rDNA DSBs via HR. Using CRISPR-based targeted rDNA DSBs, transcriptional inhibition, high-resolution imaging, and chromatin profiling, we uncover a transcription-dependent mechanism in which RNAPII, together with the DNA end-resection factor CtIP and the H3K36me3 histone marker, orchestrate the recruitment of some DNA resection factors to rDNA lesions. RNAPII inhibition disrupts nucleolar cap formation, impairs DNA end resection and induces genome instability and cell death. Mechanistically, CtIP enables RNAPII recruitment and transcriptional activation at rDNA breaks, allowing deposition of H3K36me3, a histone mark crucial for HR engagement.

Our findings reveal a previously unrecognized transcription-coupled repair program within nucleolar chromatin. We propose that a CtIP–RNAPII–H3K36me3 axis facilitates rDNA break relocation, transcriptional activation, and chromatin remodeling to enable HR at nucleolar caps. This mechanism underscores the importance of RNA-mediated genome surveillance in ribosomal DNA maintenance and highlights novel vulnerabilities that could be exploited in cancer therapy.

## Results

### Role of RNAPI, RNAPII and RNAPIII in rDNA repair

To investigate the mechanisms underlying the role of RNAPs in rDNA repair within nucleolar caps, we first selectively inhibited RNAPI, RNAPII, and RNAPIII. We took advantage of our well-characterized human sarcoma U2OS-based cell model that stably expresses the endonuclease Cas9 and a GFP-tagged version of the DDR protein NBS1 ^11, 22, 34^. Following transfection with three guide RNA vectors targeting the 5’ETS region and intergenic regions of rDNA (Fig. 1a, left), DNA repair-associated nucleolar caps formed at the nucleolar periphery within six hours (Fig.1a, right). To evaluate the impact of RNAP inhibition, cells were treated with the different inhibitors for two hours prior to fixation at the six-hour time point. Notably, the nucleolar caps associated with rDNA repair examined in this study differ from those induced by RNAPI inhibitors such as actinomycin D, BMH-21, and CX5461, which show complete absence of NBS1 (Fig. S1). To specifically investigate the role of RNAPs in rDNA damage repair—and to avoid analyzing nucleolar caps not associated with DNA repair—we focused exclusively on NBS1-positive nucleolar caps (Fig. S1). For subsequent experiments, we used BMH-21 to inhibit RNAPI, THZ1—a covalent CDK7 inhibitor previously shown to impair HR genome-wide—to inhibit RNAPII, and ML-60218 as an RNAPIII inhibitor. We observed that the proportion of cells displaying more than one NBS1- positive nucleolar cap was unaffected by BMH-21 or ML-6021, whereas treatment with the RNAPII inhibitor THZ1 led to a greater than two-fold reduction in nucleolar cap formation at the nucleolar periphery (Fig. 1b). THZ1 also significantly decreased both the average number and distribution of nucleolar caps per cell, an effect not observed upon either RNAPI or RNAPIII inhibition (Fig. 1c-d).

**Fig 1.**
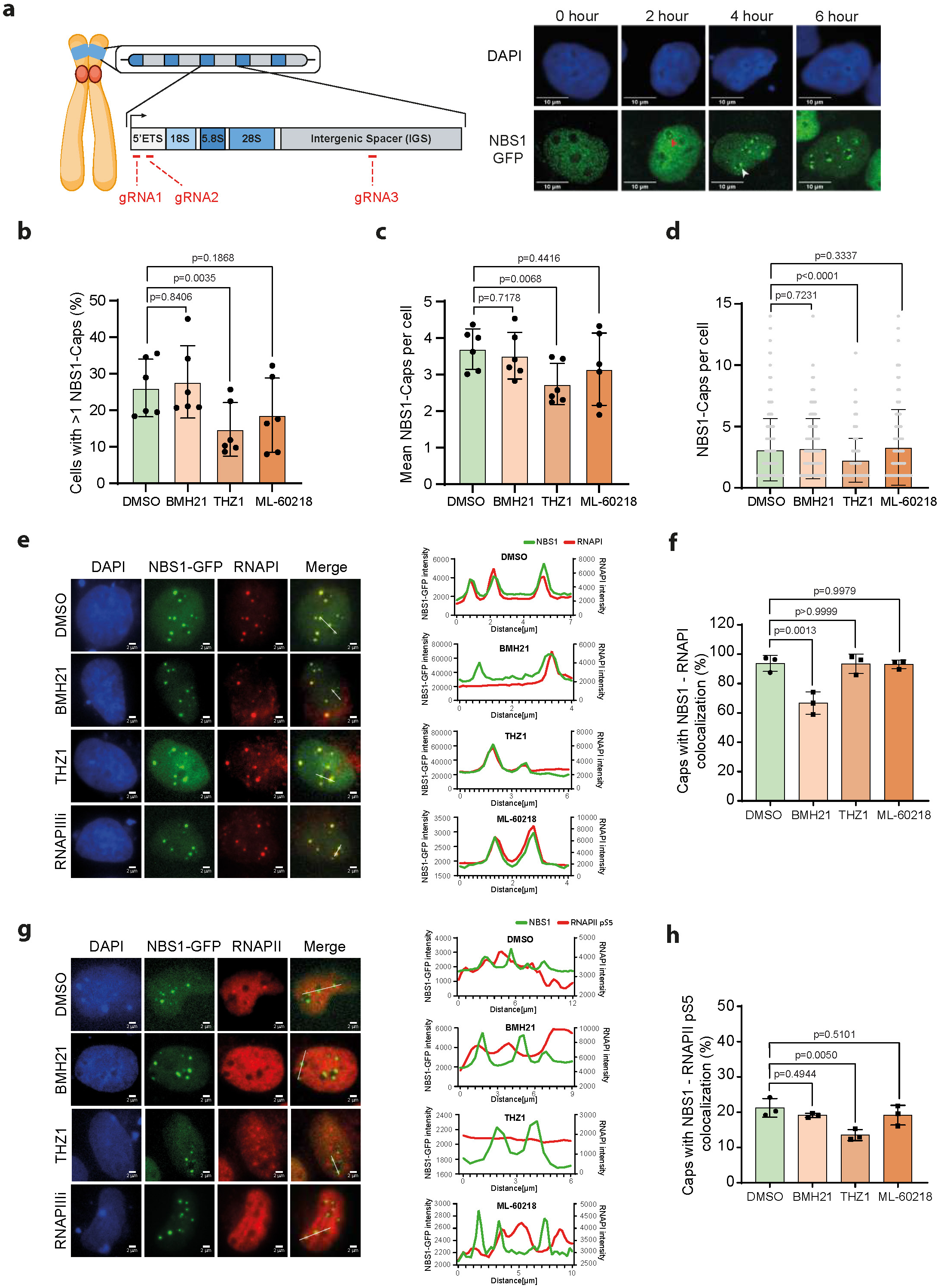

Next, we examined the presence of the three RNAPs within NBS1-positive nucleolar caps. RNAPI was detected in nearly all NBS1-positive caps. However, its recruitment was significantly reduced (by ∼35%) in BMH-21-treated cells (Fig. 1e-f), suggesting that RNAPI activity is required for robust recruitment to a subset of nucleolar caps. In contrast, neither THZ1 nor the RNAPIII inhibitor affected RNAPI localization to these caps (Fig. 1e-f).

RNAPII recruitment, assessed via phosphorylation of serine 5 on its C-terminal domain (CTD S5P)—a marker of transcriptional initiation by RNAPII—was observed in approximately 21% of NBS1-positive nucleolar caps (Fig. 1g-h). This is consistent with our previous findings suggesting that these rDNA foci may undergo repair via the HR pathway^9^ . Notably, RNAPII recruitment was not affected by inhibition of either RNAPI or RNAPIII (Fig. 1g-h). Finally, RNAPIII, detected via its RPC32 subunit, was present in only ∼8% of NBS1-positive nucleolar caps, and this fraction remained unchanged across all RNAP inhibitor treatments (Fig. S2a-b).

Together, these data indicate that while both RNAPI and RNAPII are actively involved in rDNA repair at NBS1-positive nucleolar caps, only RNAPII inhibition significantly impairs the cap formation.

### RNAPII activity is required for nucleolar cap formation upon rDNA damage

RNA transcription associated with DNA repair by HR is an essential aspect of accurate DSB repair in the nucleoplasm ^12^. Moreover, persistent rDNA breaks become relocated out of the nucleoli, thereby avoiding cis-recombination between rDNA copies to preserve the stability of these vulnerable genomic regions. We hypothesized that rDNA DSB relocation to the nucleolar periphery is also required to allow their engagement with specific DNA repair factors, including an active RNAPII for an accurate, HR-mediated DNA break repair. To test this hypothesis, we first monitored nucleolar cap formation after induction of rDNA DSBs using our U2OS-NBS1-GFP- Cas9 system transfected with gRNA vectors and treated with RNAP inhibitors. In these cells, THZ1 treatment efficiently reduced S5P of the RNAPII as revealed by nuclear immunofluorescence intensity upon staining with a RNAPII-S5P antibody (Fig. 2b-c) ^35^. Inhibition of RNAPII by THZ1 blocked initiation and elongation of nascent RNA transcripts and consequently also reduced DNA end resection as the primary step of the HR-mediated DSB repair (Fig. 2d). The reason why THZ1 treatment also partly reduces DNA resection in cells transfected by an empty vector instead of gRNA reflects the fact that RNAPII also plays a role in repair of endogenous DNA damage, as we demonstrated previously ^12^. Monitoring formation of NBS1-GFP-containing nucleolar caps after transfection of rDNA-specific gRNAs demonstrated that >25% of cells accumulated these structures in the nucleolar periphery by 6 hours after gRNA transfection (Fig. 2e-f). Treatment with THZ1 for 2 hours before the 6-hour cap formation assessment time-point (Fig. 2a) revealed an almost 3-fold decrease in the percentage of cells with NBS1 positive caps (Fig. 2e-f). We also confirmed that RNAPII inhibition impaired the number of NSB1-GFP-positive caps formed per cell, consistent with the requirement of active RNA transcription for efficient cap formation (Fig. 2g). In control experiments, we excluded the possibility that Cas9 levels were affected by RNAPII transcription inhibition during the 2 hours of treatment with THZ1 (Fig. 2h).

**Fig 2.**
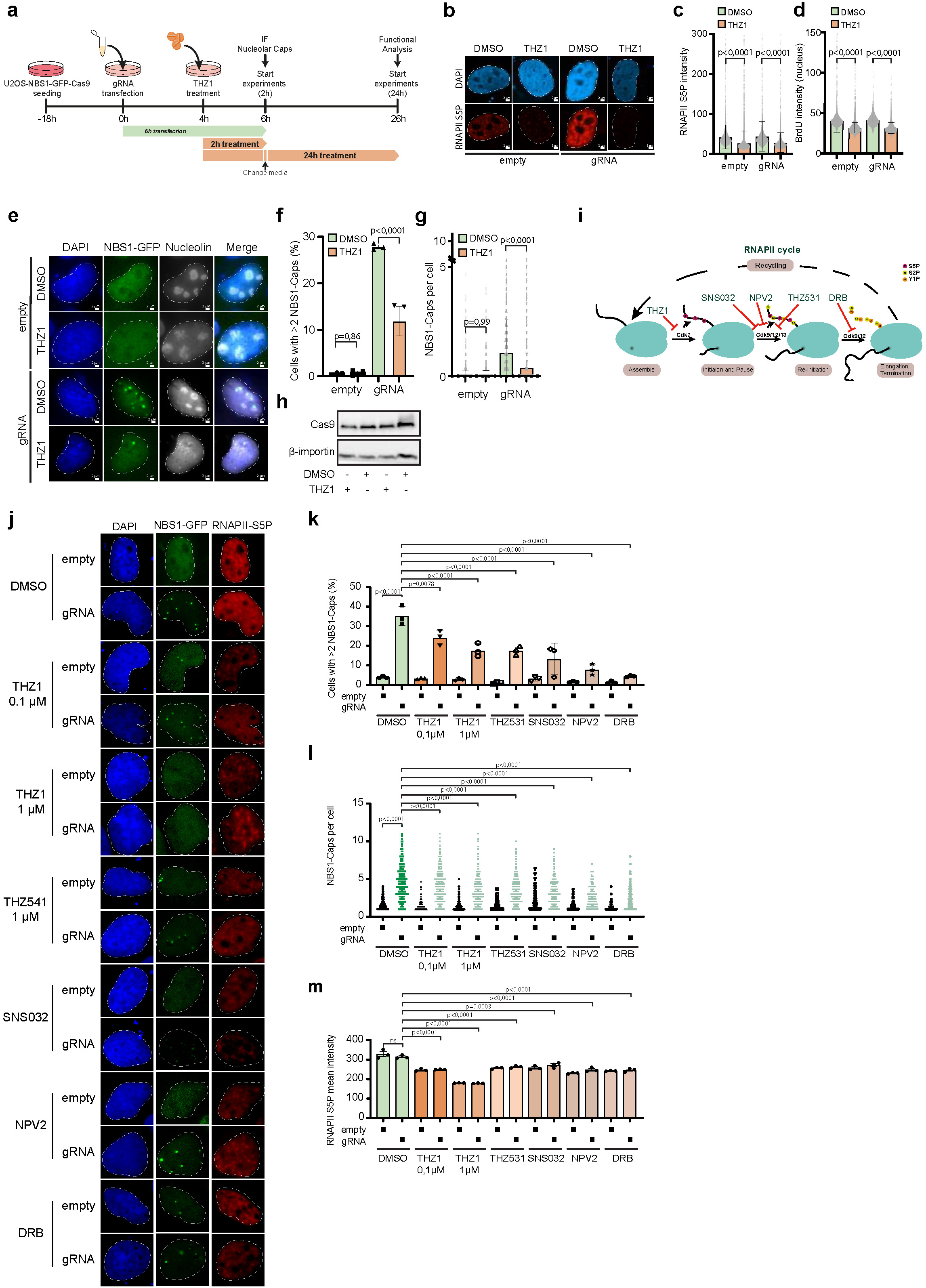

To corroborate the results obtained using stable Cas9 expression in U2OS, we further assayed rDNA DSB behavior in U2OS cells upon transient gRNA transfection and direct transfer of ectopic recombinant Cas9 protein, thereby avoiding the plasmid integration events and potential off-target effects due to stable Cas9 maintenance in the cells. Nucleolar caps formed more rapidly using this transient expression approach compared with the plasmid-based expression system ^6^. Indeed, the cells showed nucleolar cap formation already during the first 3 hours after gRNA/Cas9 transfection (Fig. S3a). THZ1 treatment immediately after transfection reduced nucleolar cap formation at 1 hour after transfection and even more robustly, up to 4-fold, after 3 hours of THZ1 exposure (Fig. S3a. This approach excludes any possible negative impact of either Cas9 or NBS1-GFP stable expression during THZ1 treatment (Fig. S3b), which might affect S5P phosphorylation of the RNAPII (Fig. S3c).

The decrease of RNA transcription initiation upon THZ1 treatment was accompanied by a defective DNA end resection as the primary step of the HR-mediated DSB repair (Fig. 2d), consistent with the dependence of HR repair on nascent RNA synthesis. Taken together, our data suggest that RNAPII inhibition by THZ1 impairs nucleolar cap formation, probably due to the requirement of nascent RNA for efficient assembly, and hence function, of the HR machinery at the nucleolar periphery.

### Transcription initiation and elongation promote accurate rDNA DSB repair

While THZ1 is a potent specific inhibitor of CDK7, hence blocking the initiation of nascent RNA transcript, we asked if other RNAPII inhibitors targeting CDK7, CDK9, CDK12 and CDK13 would also impair cap formation upon induction of rDNA breaks (Fig. 2i). First, even a 10-fold lower concentration of THZ1 (0.1 μM), reduced the percentage of cells with nucleolar caps and number of caps per cells upon Cas9-induced rDNA DSBs (Fig. 2j-l). THZ531 is a covalent inhibitor of CDK12/CDK13 when used at low concentrations, 158 and 69 nM, respectively, but it can affect CDK7 and CDK9 at high concentrations (>8.5 μM). THZ531 used at a low concentration (1 μM) had a similar effect to that caused by THZ1 in NBS1-GFP-Cas9 cells after gRNA-vectors transfection to generate rDNA breaks, namely a 2-fold decrease of nucleolar cap formation (Fig. 2j-l). This effect suggests that transcription elongation promoted by the CDK12/13 kinase-mediated serine 2 phosphorylation of the RNAPII CTD may also contribute to nucleolar cap formation in cells with rDNA DSBs. However, we noticed that THZ531 at this concentration also decreased S5P of the RNAPII CTD domain, similarly to THZ1 (Fig. 2m), suggesting that mainly initiation and, possibly, elongation steps of RNAPII transcription promote cap accumulation. Then, we investigated other CDK9-targeting drugs that operate through different mechanisms. THAL-SNS- 032 is a selective CDK9 PROTAC degrader consisting of a CDK-binding SNS-032 ligand linked to a thalidomide derivative that binds the E3 ubiquitin ligase Cereblon (CRBN)^36^. NPV-2 is a potent and selective ATP-competitive inhibitor of CDK9^37^ that inhibits RNAPII at low concentrations. Treatment with either of these inhibitors during 2-3 hours before nucleolar cap assessment upon rDNA DSB induction demonstrated that any RNAPII dysfunction in S5P-CTD caused by CDK9 inhibition impairs rDNA repair associated with both a decreasing percentage of cells with NBS1- marked nucleolar caps, and the number of caps per cells (Fig. 2j-l). These data indicate that CDK9 inhibition causes a deficiency in rDNA repair *via* impairment of RNA transcription. Finally, we used DRB, a classic inhibitor of transcription elongation by RNAPII, yet with a broader spectrum of targets than the other inhibitors used above. DRB treatment also resulted in a lower percentage of cells with NBS1-foci and NBS1-cap structures relative to control DMSO-treated cells (Fig. 2j- l). To minimize the likelihood of any off-target effect of THZ1, we used another known RNAPII inhibitor, triptolide, that affects RNA transcription initiation by RNAPII degradation via proteasome ^12, 38^. We observed similar nucleolar cap formation deficiency after triptolide treatment, demonstrating that the inhibition of transcription initiation by RNAPII indeed compromises nucleolar cap formation (Fig. S3d-e). All the RNAPII inhibitory compounds used in this study consistently showed that nascent RNA transcription by RNAPII is essential for cap formation measured by NBS1 accumulation.

### RNAPII colocalizes with nucleolar caps upon rDNA DSB formation

Our recent studies showed that transcription initiation promotes HR factor recruitment at whole genome DNA DSBs in nucleoplasm ^12, 23, 26, 39^. To further study the relevance of the RNAPII function for rDNA BSB repair, we next wondered if the transcription complexes are recruited actively to the nucleolar caps. We utilized high-resolution fluorescence imaging of NBS1-GFP- Cas9 U2OS cells to assess the colocalization of additional factors at nucleolar caps. First, cells were co-stained with antibodies to transcription initiation markers such as S5P residue in the CTD domain of the RNAPII. As shown in Fig. 3a the RNAPII-S5P staining is pan-nuclear but reveals accumulation in distinct foci close to the nucleolar periphery. To assess whether the transcription initiation complex is located in the nucleolar caps, we analyzed the dynamic profile of rDNA DSBs marked by NBS1-GFP, along with the transcription initiation marker RNAPII-S5P. We observed RNAPII-S5P recruitment to nucleolar caps, significantly colocalizing with NBS1, suggesting that active nascent transcription in these regions is spatially coordinated with rDNA repair (Fig. 3a). Under these conditions, we also detected a later stage of RNA nascent process using S2P of the CTD domains that drives transcription elongation (RNAPII-S2P). RNAPII-S2P remained localised at the nucleolar caps, although it showed a diffuse colocalization pattern in some of them (Fig. 3b). These results suggest that early transcription steps do occur in the vicinity of the rDNA DSBs at the periphery of the nucleolus, consistent with a functional link between active RNAPII and rDNA repair^12^. Finally, we analysed the profile for other phosphorylations in the CTD of the RNAPII, specifically in Tyrosine 1 (RNAPII-Y1P), which some studies have related to DSB repair ^25^. Our colocalization analysis showed that RNAPII-Y1P occupancy in the nucleolar caps was limited to the periphery of the rDNA DSB-containing regions marked by NBS1-GFP (Fig. 3c), suggesting that this modification *per se* is unlikely to be directly required for the rDNA repair, despite it is locally connected with the area of RNA-coupled rDNA repair. Taken together, these results suggest a relationship between nascent RNA transcription by RNAPII and nucleolar caps.

**Fig 3.**
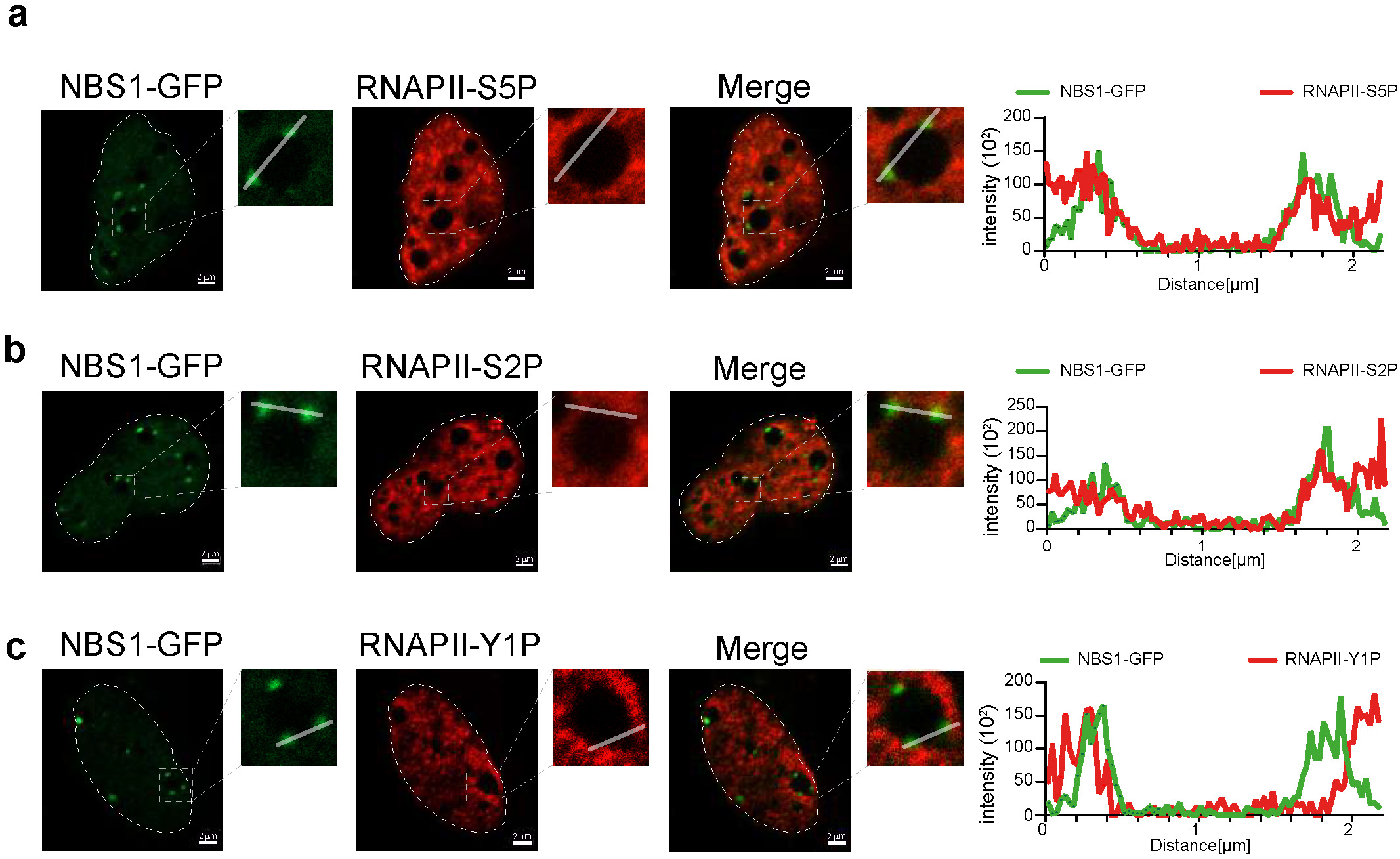

As we mentioned previously, RNAPIII has recently been associated with promoting HR of DSBs, similarly to RNAPII^23^. Therefore, we analyzed RPC32, an RNAPIII-specific subunit, in further detail. High resolution images showed that RNAPIII does not colocalize completely with NBS1- GFP in the nucleolar caps, but partial colocalization was detected (Fig. S2c) Notably, a subset of nucleolar caps showed a very robust match between NBS1 and RPC32 proteins while other nucleolar caps lacked RPC32 entirely, even in the same cell (Fig. S2c). These results suggest that RNAPII is highly enriched in nucleolar caps, whereas RNAPIII is only partially present and likely plays a more limited, context-dependent role in this process.

### RNAPII inhibition impairs recruitment of HR factors to nucleolar caps

The relocation of rDNA DSBs from the nucleolar interior to anchoring points at the nucleolar periphery is thought to render rDNA breaks accessible to repair factors that are excluded from the nucleoli ^18^. To gain mechanistic insight into this process, we examined whether DNA resection following CRISPR-induced rDNA DSBs occurs within nucleolar caps and whether this is influenced by RNAPII inhibition. We assessed CtIP localization at NBS1-positive nucleolar caps to determine whether the observed colocalization pattern depends on ongoing transcription. CtIP was detected in over 60% of repair-associated nucleolar caps (Fig. 4a-b), However, treatment with transcription inhibitors such as THZ1, DRB, and Triptolide reduced CtIP recruitment by more than 20% (Fig. 4a–b). To confirm these results, we carried out a Proximity Ligation Assay (PLA) to assess recruitment of CtIP after treatment with the RNAPII inhibitor, THZ1. Ribosomal DSBs induction showed interaction between NBS1 and CtIP partially dependent of RNAPII transcription, as suggested by lowered PLA spots after THZ1 treatment (Fig. 4c). Treatment with the RNAPII inhibitor did not alter the cellular CtIP protein level, thereby excluding the possible overall CtIP deficiency under these conditions (Fig. S4a). To rule out the possibility that THZ1 affects other DSB repair factors such as 53BP1, we analyzed its colocalization with NBS1 under the same conditions and found it to be unaffected by RNAPII inhibition (Fig. S4b-c).

**Fig 4.**
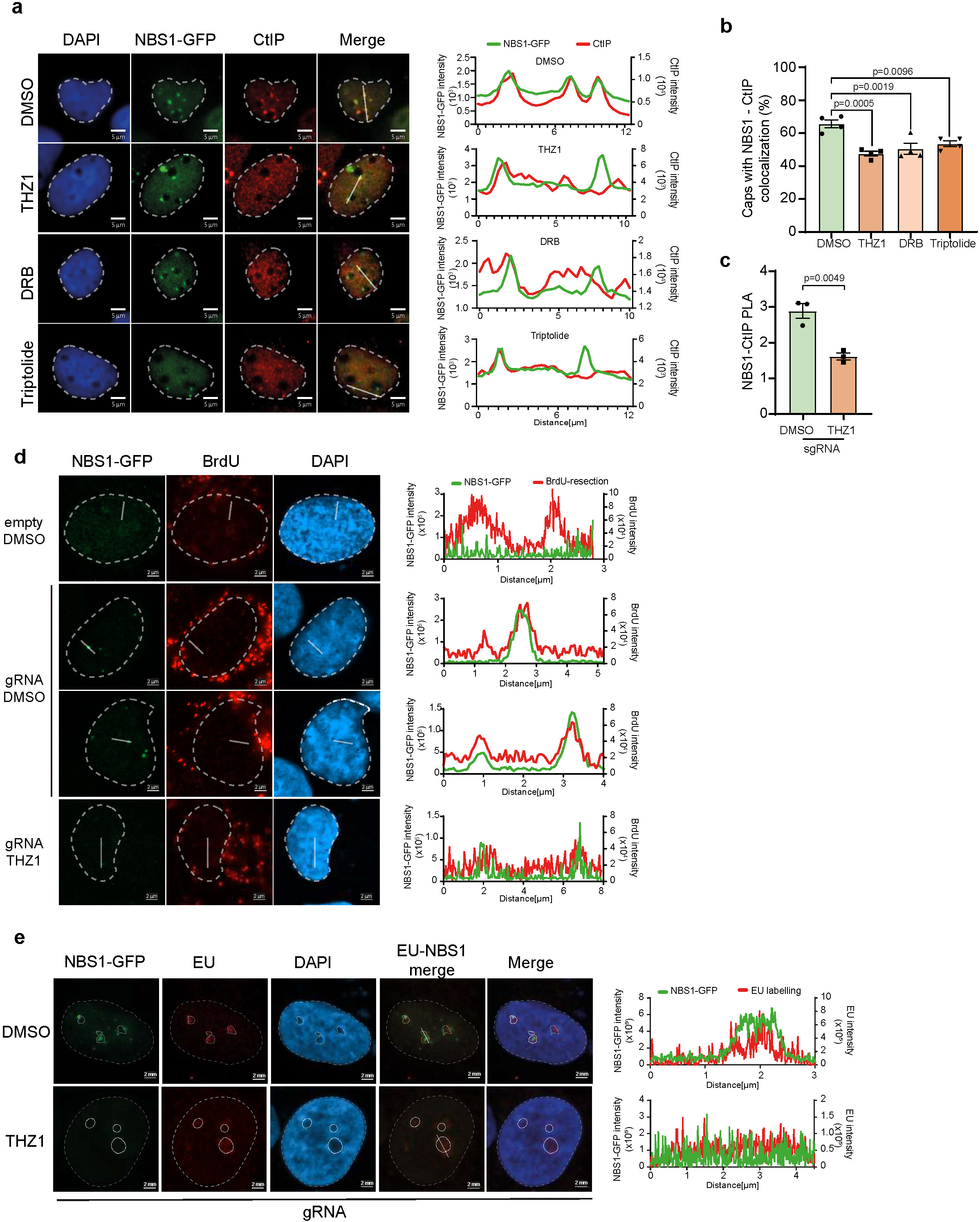

To further evaluate the impact of rDNA resection upon RNAPII inhibition, we detected the ssDNA stretches formed by rDNA end resection. To do so, we employed immunostaining for native BrdU incorporation, reflecting accessibility of the resected 5’ DNA strand, an ssDNA intermediate in the HR repair mechanism ^12^. Profile analysis showed colocalization peaks for the NBS1-GFP and BrdU staining, indicating an active DNA resection process inside the analysed nucleolar caps (Fig. 4d). We wanted to investigate whether this decrease is likely due to the requirement for RNAPII activity in the nucleolar caps, in order to repair the rDNA breaks by the HR pathway. To validate the activity of RNAPII within the nucleolar caps, we conducted an analysis of nascent RNA incorporation using EU. This analysis provided evidence that RNA transcription occurs inside the nucleolar caps and is necessary for the formation of these rDNA DSB repair structures, as indicated by the colocalization pick of nascent RNA with DNA repair factors such as NBS1 (Fig. 4e). Altogether, our data indicate that the nascent RNA plays a crucial role in recruiting the HR repair machinery in the rDNA chromatin context.

### Ribosome biogenesis remains unaffected by rDNA Double-Strand Breaks

We demonstrated that RNAPII inhibition under conditions of rDNA DSB generation by CRISPR technology impaired the rDNA damage repair. We therefore wondered whether the treatment with THZ1 might have negatively affected ribosome biogenesis and whether such potential effect would be enhanced upon generation of rDNA DSBs in our model system. To test so, we performed polysome profile analysis. Our results indicate that neither ribosome biogenesis nor translation were significantly affected by the THZ1 treatment, the induction of rDNA double-strand breaks (DSBs), or the combination of both, within 6 hours after gRNA transfection (Fig. 5a, left). At the 24-hours post rDNA DSB induction, the observed polysome profiles primarily reflected the impact of RNAPII inhibition—namely, mRNA depletion—yet, once again, rDNA DSB induction does not appear to disrupt translational activity (Fig. 5a, right). To further analyze the potential consequences of THZ1 treatment for ribosomal biogenesis at 24 hours post rDNA damage induction, we analysed pre-rRNA processing in U2OS cells by assessing the levels of pre- and mature rRNAs by northern blotting (Fig. 5b). As shown in Fig. 5c-d, inhibition of RNAPII significantly affected pre-rRNA processing. Thus, upon THZ1 treatment for 24 hours, an accumulation of early (e.g. 47S/45S pre- rRNAs) and later (e.g. 30S, 32S, 12S pre-rRNAs) pre-rRNAs was observed (Fig. 5c-d). In contrast, steady-state levels of mature 18S and 28S rRNAs remained unaffected upon the treatment, likely reflecting the stable nature of preexisting ribosomes. The functional significance of these pre-rRNA processing alterations is unclear. However, it could be due to limited levels of specific protein assembly factors and/or their function following RNAPII inhibition. Importantly, formation of the CRISPR-induced rDNA DSBs alone neither impeded pre-rRNA transcription nor enhanced the observed pre-rRNA processing defects induced by THZ1 treatment, indicating that this type of rDNA damage did not reduce the overall number of actively transcribed rDNA copies, a result that was also consistent with the polysome profile data (Fig. 5a). These results indicated that, *per se*, the rDNA damage induced in our model system does not cause any major defects of ribosome biogenesis and function that could influence the interpretation of our results (see Discussion).

**Fig 5.**
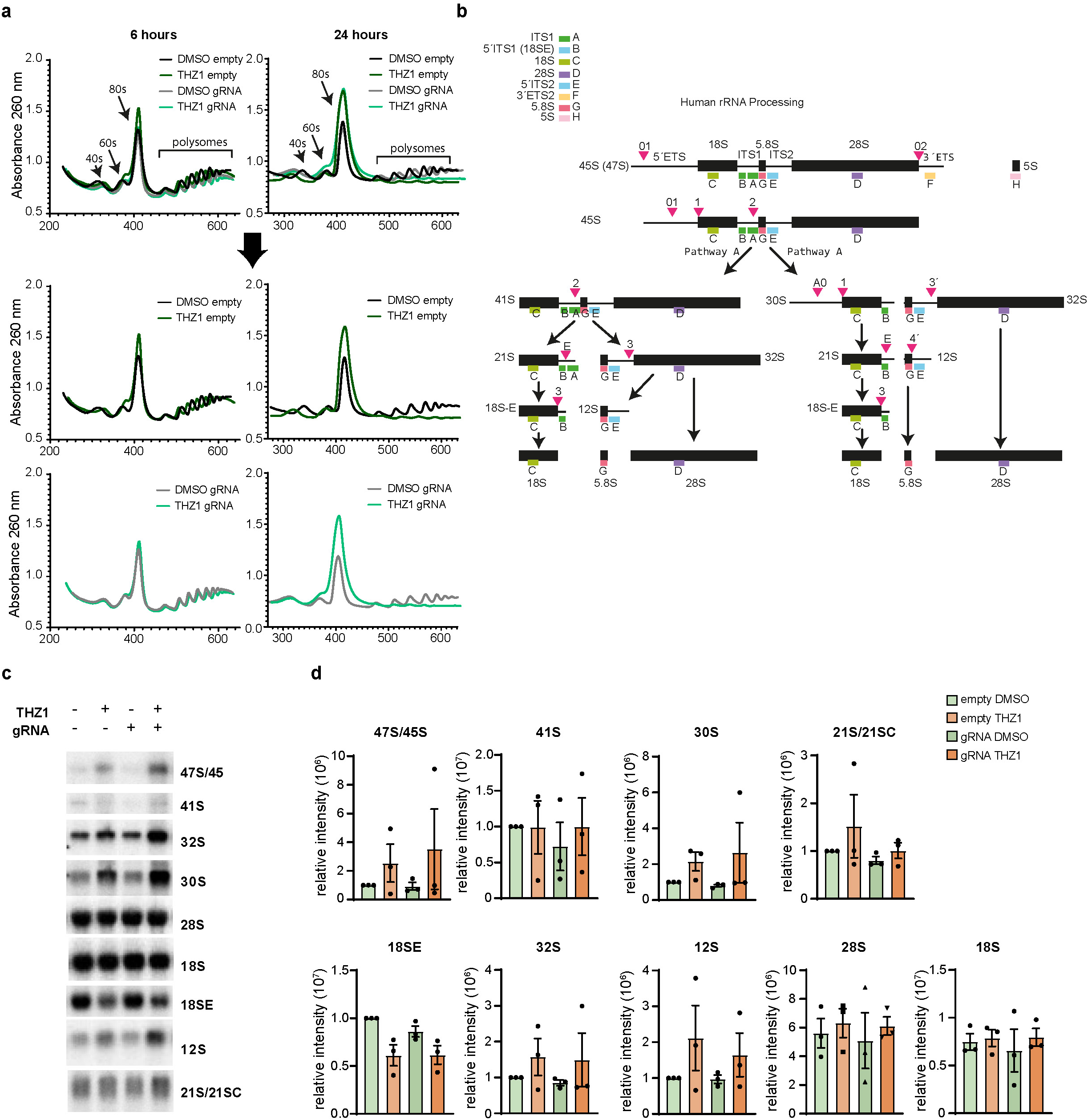

### CtIP regulates RNAPII recruitment for rDNA repair

To provide more mechanistic insights into the role of RNAPII in nucleolar caps-associated HR of rDNA breakage, we next investigated whether CtIP may facilitate RNAPII recruitment^9^ to the rDNA DSBs in our model. Therefore, we knocked down CtIP to determine whether it is similarly required for rDNA repair—especially considering that both RNAPII and CtIP are mostly present in the nucleoplasm. Indeed, silencing CtIP impaired RNAPII recruitment to nucleolar caps following rDNA DSB induction (Fig. 6a-c). Moreover, BRCA1 accumulation in nucleolar caps was also markedly reduced in CtIP-depleted cells, suggesting that CtIP is essential for the recruitment of the DNA resection machinery at these specific sites (Fig. 6d-e). In contrast, localization of the NHEJ factor 53BP1 within nucleolar caps remained unaffected by CtIP silencing and was observed in the majority of nucleolar caps following rDNA damage (Fig. 6f-g). These findings underscore the emerging role of the CtIP–RNAPII axis in HR-mediated repair of rDNA DSBs within nucleolar caps, while highlighting the distinct and CtIP-independent recruitment of NHEJ factors such as 53BP1.

**Fig 6.**
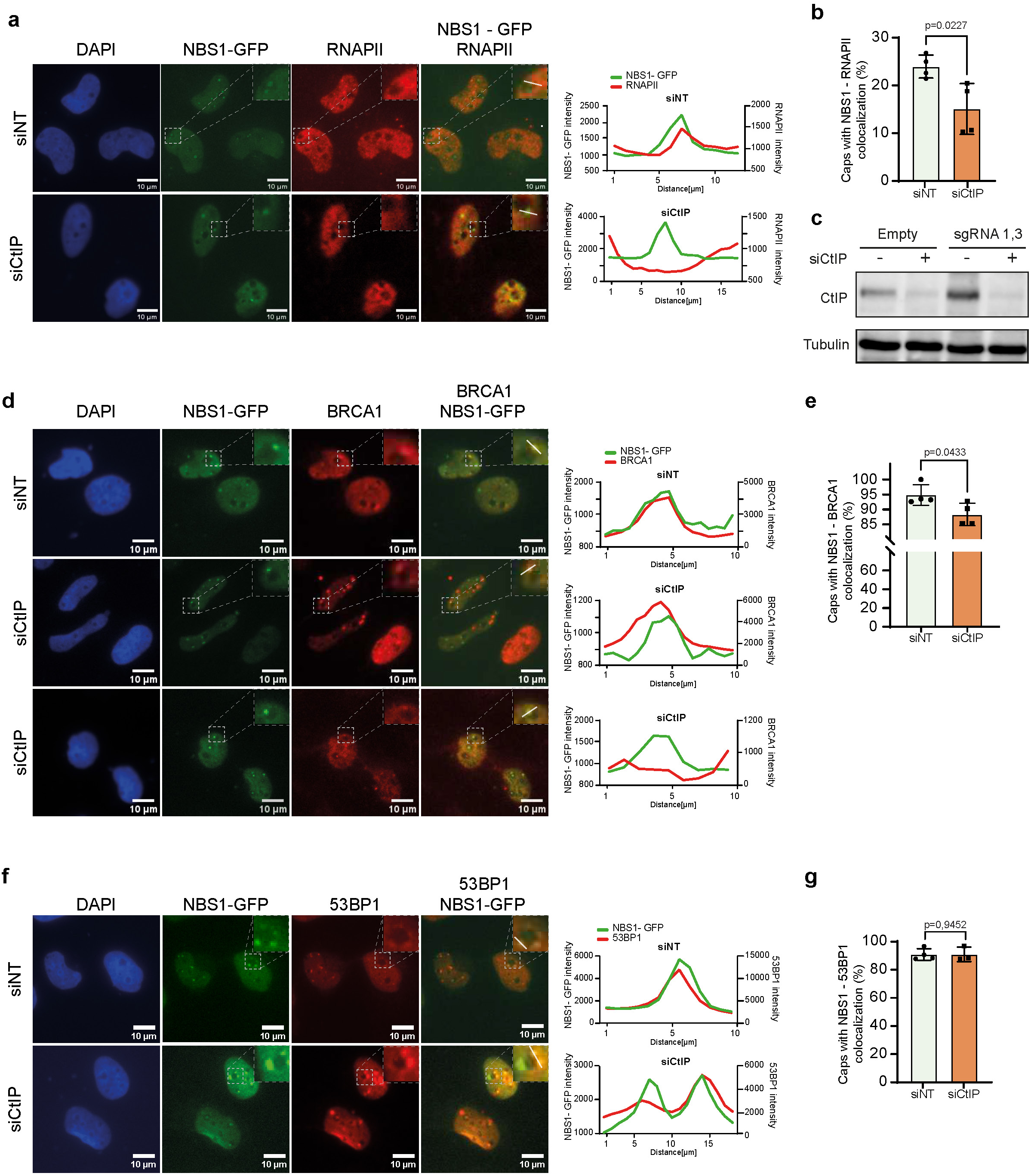

To investigate whether CtIP contributes to RNAPII-dependent transcription at damaged rDNA sites, we assessed levels of H3K36 trimethylation (H3K36me3), a histone mark classically associated with RNAPII transcription elongation and with the recruitment of DNA repair factors at DSBs ^40, 41^. Silencing CtIP led to a pronounced reduction in the H3K36me3 immunostaining signal within nucleolar caps (Fig. 7a-b) and a two-fold decrease in the spatial coordination between RNAPII and H3K36me3 at sites of rDNA repair, as measured by colocalization analysis (Fig. 7c). The CtIP/RNAPII deficiency impaired also BRCA1 recruitment to rDNA breaks (Fig. 7d-e), suggesting that H3K36me3 may serve as a critical chromatin signal that facilitates HR factor recruitment and rDNA relocalization during DSB repair. Importantly, we observed an increase in nucleolar-associated H3K36me3 intensity in response to rDNA DSBs induced by CRISPR-based targeting (Fig. S5), supporting a mechanistic model in which co-transcriptional repair of rDNA breaks is coordinated by RNAPII and DNA resection machinery, and regulated through H3K36me3-dependent chromatin remodelling (Fig. 7f).

**Fig 7.**
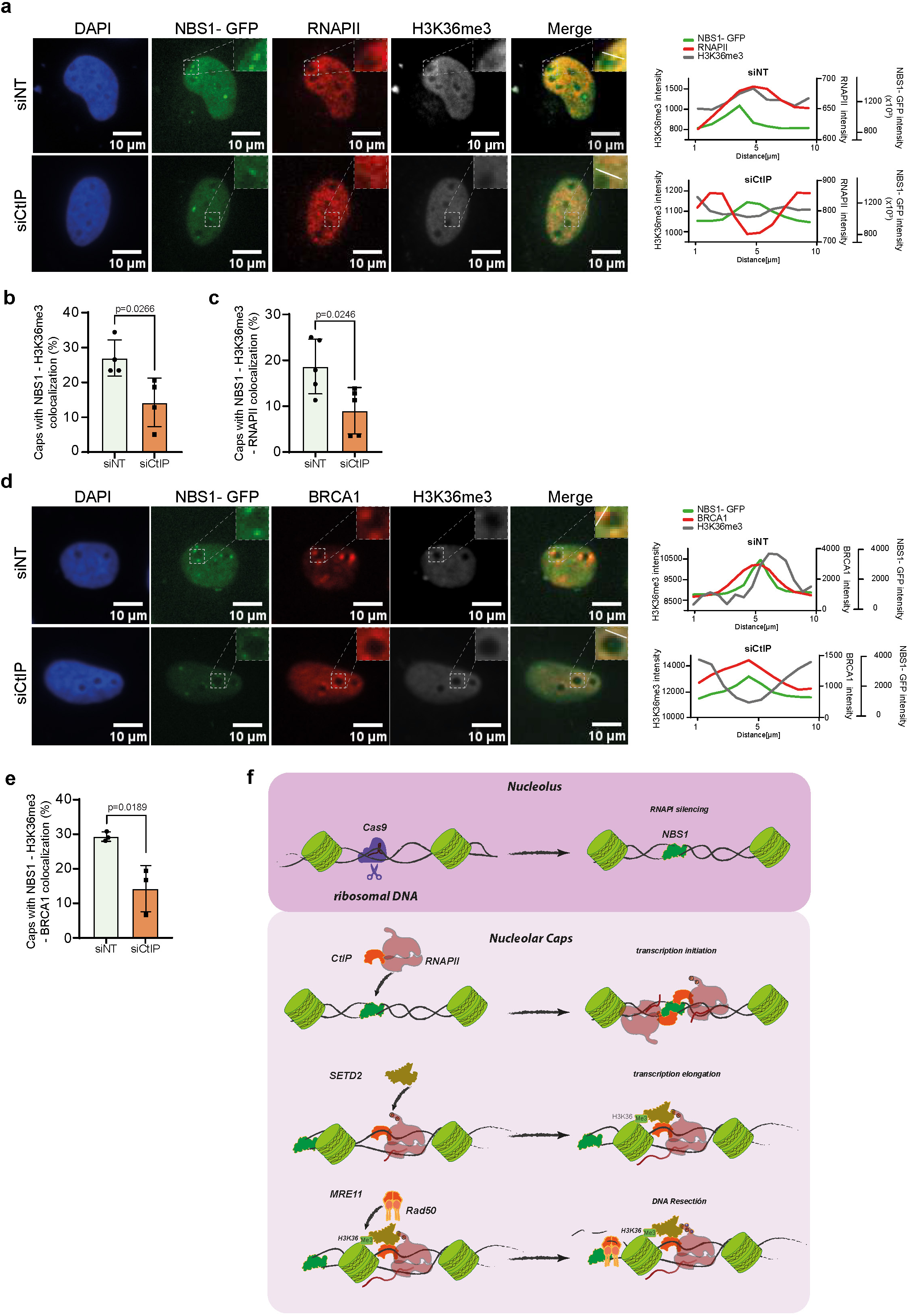

### RNAPII inhibition provokes rDNA instability and cell death in cancer cells

To assess the biological longer-term consequences of rDNA damage upon RNAPII inhibition, we first examined cell survival by clonogenic assay after rDNA DSB induction and THZ1 treatment. Induction of rDNA DSBs resulted in a robust 3-fold reduction of cell colony counts (Fig. 8a), consistent with previous work ^42^. Furthermore, THZ1 treatment alone sensitized the undamaged tumor cells, likely because RNAPII inhibition causes endogenous whole genome DNA damage^12^. We observed an increased, albeit not synergistic effect on reducing colony formation when combining the rDNA DSB induction with RNAPII inhibition (Fig. 8a and S6a). Additive effects of targeting components of the same pathway can occur, here when rDNA damage is combined with RNAPII inhibitor THZ1, due to the requirement to repair the endogenous DSBs outside rDNA, thereby achieving a stronger blockade of essential processes and ultimately amplifying cell growth inhibition or cell death. To examine this phenotype further, we performed an MTT assay based on the reduction of the tetrazolium salt MTT to a dark-coloured formazan product by metabolically active cells, to obtain complementary assessment of cell viability. The damage in rDNA alone reduced (2-fold) cell viability (Fig. 8b). Likewise, RNAPII inhibition by THZ1 negatively impacted the cell viability of both rDNA-damaged and undamaged cells, with a tendency for an additive effect when rDNA damage and THZ1 treatment were combined (Fig. 8b). The results were similar but more pronounced after 24 hours of THZ1 treatment (Fig. S6b).

**Fig 8.**
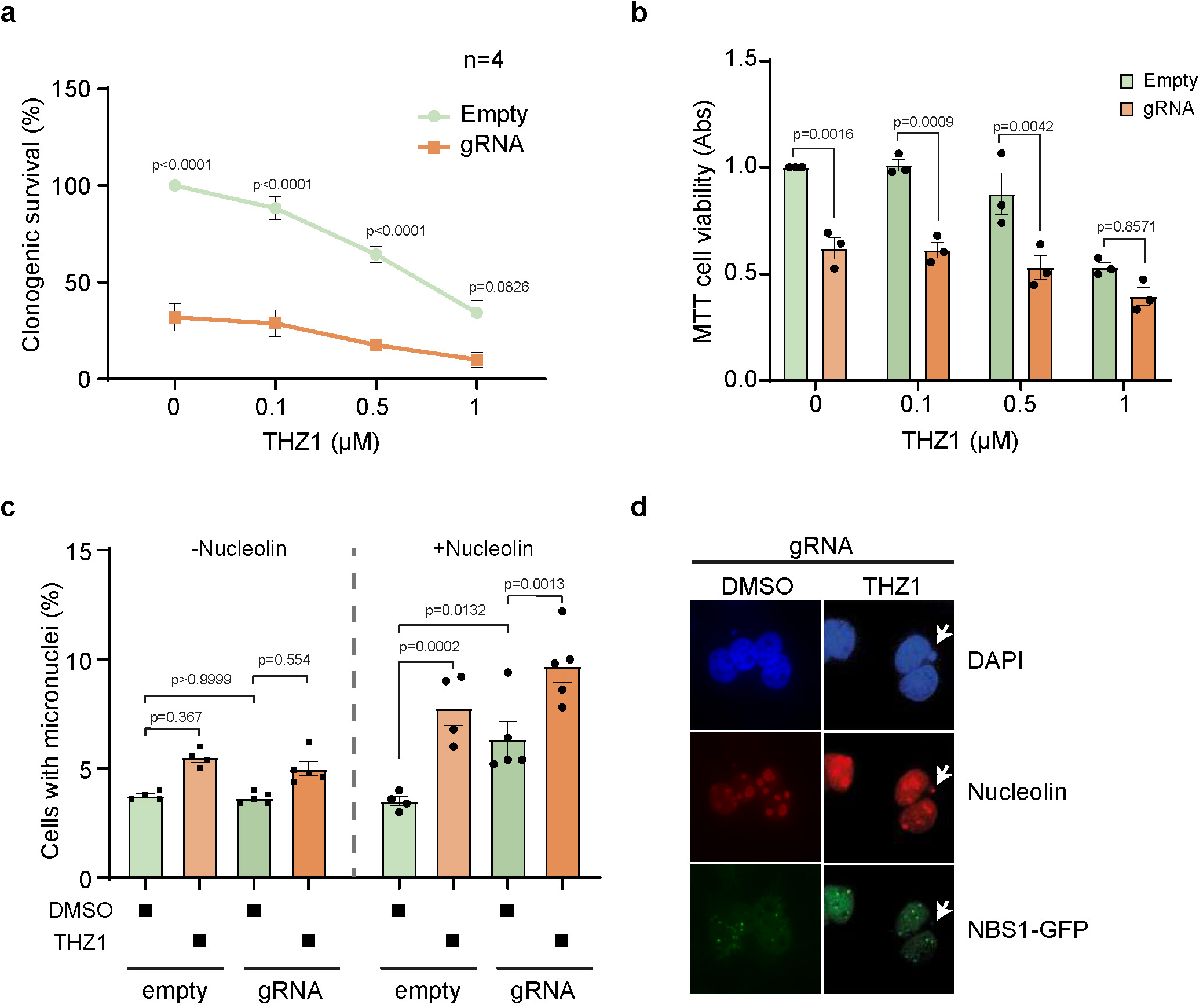

It is known that nuclear DNA damage can cause mitochondrial dysfunction, and changes in mitochondrial membrane potential may undermine cell viability though apoptosis ^43^. Thus, we analysed mitochondrial membrane potential after THZ1 treatment for 2 hours in rDNA-damaged and undamaged cells, but no significant differences were observed (Fig. S6c-d). There was a modest, non-significant decrease in membrane potential in cells treated with THZ1 for 24 hours (Fig. S6e). As THZ1 impairs nucleolar cap formation, and defective DNA repair usually leads to an increase in genomic instability, we next investigated whether RNAPII inhibition after rDNA DSB induction may impact genome stability. For this purpose, we analysed micronuclei formation frequency, one of the indicators of genome instability due to DNA damage ^44^. After rDNA DSB induction, a 2-fold increase of micronuclei formation was observed (Fig. 8c), further enhanced by RNAPII inhibition upon THZ1 treatment (Fig. 8c-d). Notably, rDNA regions were present in the formed micronuclei after rDNA lesion induction and transcription inhibition conditions (Fig. 8d). After 24 hours of THZ1 treatment, the micronuclei formation results were similar (Fig. S6f). Altogether, our results emphasise the importance of rDNA repair under genotoxic nucleolar stress and highlight the important role of RNAPII in the maintenance of nucleolar homeostasis and rDNA stability.

## Discussion

Ribosomal DNA represents one of the most transcriptionally active and repetitive genomic regions, inherently prone to instability under replicative or transcriptional stress, a condition frequently amplified in cancer cells due to deregulated ribosome biogenesis and elevated protein synthesis ^1–3^. While being transcriptionally dominated by RNAPI, rDNA is also susceptible to DNA DSBs, particularly at the intersection of replication and transcription machinery ^6,7^. The rDNA lesions pose a unique challenge due to the repetitive nature of rDNA, requiring specialized repair pathways to prevent catastrophic chromosomal rearrangements ^8,14,35^.

Although RNAPII was not previously thought to function in the nucleolus, it has recently been shown to regulate RNAPI transcription and, under genotoxic stress, driving transcription of the rDNA intergenic spacer to sequester aberrant transcripts for DNA repair ^31, 32^. However, whether RNAPII contributes to the regulation of the specialized nucleolar nDDR remains unclear, and in this work, we extend its role by demonstrating its relevance in maintaining genomic stability within this region. Our findings demonstrate an active presence of RNAPII to rDNA regions that are not transcribed by RNAPI, suggesting a functional role beyond passive association. This support previous data about the role of RNAPII in DNA resection on untranscribed regions ^12, 26, 27^, and not just in active region ^21, 45^, including rDNA genes. Furthermore, our data aligns with emerging evidence supporting a complementary role for RNAPII within the nucleolus alongside RNAPI ^31,32^.

This process is orchestrated via a transcription-coupled mechanism involving RNAPII recruitment to nucleolar caps, perinucleolar structures that serve as peripheral subdomains for active DNA repair ^20, 34, 46^. Pharmacological inhibition of RNAPII by CDK7 (THZ1), CDK9, CDK12/13 inhibitors, as well as elongation blockers (DRB, triptolide), significantly impaired nucleolar cap formation and HR factor assembly, highlighting a transcription-dependent switch for pathway engagement. These findings extend earlier work that identified RNAPII as a determinant of repair pathway choice at nucloplasmic DSBs ^4,9,20^, now applied to the nucleolar context.

Mechanistically, we show that transcription initiation, and likely also elongation, are necessary for efficient recruitment of RNAPII to rDNA damage foci, as reflected by Ser5 and Ser2 phosphorylation of its C-terminal domain (CTD), respectively. Pharmacological inhibition of these critical regulatory phosphorylations of RNAPII abrogates the early assembly of HR components. The co-localization of NBS1 with markers of transcriptionally active RNAPII confirms that rDNA lesions are transcriptionally licensed for repair, consistent with previous models where nascent RNA guides HR machinery to nucleoplasmic DSBs *via* RNA–protein scaffolds. ^9,20,26,27^ Furthermore, our high-resolution imaging and biochemical data confirm that CtIP acts as a master regulator in this rDNA repair mechanism. Knock-down of CtIP not only prevented RNAPII recruitment but also reduced H3K36me3 levels at nucleolar caps, a chromatin modification catalyzed by SETD2 and essential for BRCA1 loading and HR progression ^33,34,42^. The latter result underscores a previously uncharacterized role for CtIP in controlling transcription-coupled chromatin remodeling in the nucleolus, acting upstream of RNAPII-SETD2 in rDNA repair. These data are consistent with reports positioning CtIP as a cofactor for DNA end resection, acting in concert with the MRN complex and CDK-dependent phosphorylation to license HR ^39–41^. Our data also showed that the CtIP-dependent reduction in H3K36me3 levels following rDNA damage has significant implications for rDNA repair pathway choice, highlighting the role of this histone mark in chromatin remodeling specifically at rDNA loci. These results therefore establish a transcription– chromatin modification axis that is both spatially and mechanistically linked to HR engagement at ribosomal chromatin. The increase in H3K36me3 levels upon CRISPR-induced rDNA DSBs further strengthens this model, suggesting that damage-induced modulation of certain chromatin states facilitate rDNA repair.

Interestingly, the mechanism elucidated in our present study appears to be exclusive to RNAPII, as inhibition of either RNAPI and or RNAPIII did not impact nucleolar cap formation. This is in line with recent data positioning RNAPIII as a context-dependent factor in DSB repair ^4^, and with the known role of RNAPII in producing nascent transcripts that scaffold HR complexes at DNA breaks ^9,20,26^. Although RNAPIII components were occasionally detected in nucleolar caps, their inconsistent localization suggests a supportive, rather than central, role in rDNA repair—potentially influenced by local chromatin environments or transcriptional states ^4,18^.

Our data also support a model in which persistent rDNA DSBs that are not repaired by NHEJ inside the nucleoli become selectively relocated to nucleolar caps for repair, reinforcing prior reports that nucleolar stress triggers spatial segregation of damaged rDNA to the nucleolar periphery^47–49^ ^15–17,21^. Inhibiting RNAPII function not only impairs such rDNA relocation but also reduces the recruitment of early repair factors such as CtIP. This is consistent with prior work highlighting the interdependence of transcription and DNA end resection in HR activation.^9,20,40,41^

Importantly, we show that RNAPII inhibition compromises genome stability and cell viability, particularly in the context of rDNA damage. Increased micronuclei formation following combined RNAPII inhibition and rDNA DSB induction highlights the genomic consequences of defective rDNA repair^44^ .Such micronuclei frequently contained rDNA, indicating that unresolved breaks in ribosomal chromatin contribute to chromosomal fragmentation and mitotic errors—phenomena tightly associated with tumorigenesis ^30,32,36^.

Despite the central role of rDNA in ribosome biogenesis, we show that rDNA instability does not necessarily translate into immediate disruption of ribosomal function. Indeed, our data reveal that CRISPR-induced rDNA DSBs do not perturb rRNA transcription, processing, or polysome formation within short-term experiments. Only when RNAPII activity is inhibited we observed defects in rRNA maturation and translation, consistent with the broader role of RNAPII in transcriptional regulation of ribosome assembly factors and non-coding RNAs, including snoRNAs ^43,49^. These observations indicate that the genome maintenance function of RNAPII in rDNA repair is separable from its canonical transcriptional roles—at least over acute timescales.

Together, our findings define a previously unrecognized co-transcriptional mechanism that safeguards ribosomal genome integrity. We propose a model in which CtIP enables RNAPII access to damaged rDNA regions, promoting H3K36me3 deposition and BRCA1 loading within nucleolar caps. This orchestrated response coordinates spatial repositioning of rDNA breaks with transcriptional and chromatin-based repair signals. The broader implication of our present data is that transcription-coupled HR-mediated DNA DSB repair extends beyond the coding sequences in gene bodies and includes also repetitive, non-coding genomic compartments such as the nucleolus, thereby redefining our understanding of the genome surveillance landscape.

## Materials and Methods

### Cell culture

U2O-Cas9-NBS1-GFP cells ^34^ were cultured in Dulbecco’s Modified Eagle Medium (DMEM) with high glucose plus GlutaMax supplemented with 10% fetal bovine serum (FBS; Biowest), 100 U/ml penicillin and 100 μg/ml streptomycin (Biowest) at 37 °C in 5% CO2. U2O-Cas9-NBS1-GFP stably transfected cells were maintained by adding 1 μg/ml puromycin and 0.5 mg/ml neomycin G418 to the medium. siRNAs against CtIP (GCUAAAACAGGAACGAAUC) and a control non-targeting siRNA sequence (Sigma Aldrich) were transfected with the RNAiMax lipofectamine reagent mix (Life Technologies), according to the manufacturer’s instructions. Cells were cultured in the presence of different inhibitors (Table 1) using concentrations mentioned in each figure.

**Table 1.** RNAPs inhibitors.

| **Drug** | **Reference** | **Commercial** | **Use**<br>**Concentration** | **Time of**<br>**treatment** |
| --- | --- | --- | --- | --- |
| **THZ1** | 532372 | CalbioChem | 1 μM | 2 hours |
| **THZ531** | HY-103618- | MedChemExpress | 1 μM | 2 hours |
| **NPV2** | 1702809-17-3 | R&D systems | 250 nM | 3 hours |
| **THAL-SNS032** | 6532/5 | R&D systems | 250 nM | 3 hours |
| **DRB** | D1916 | Sigma | 100 μM | 2 hours |
| **Triptolide** | T3652 | Sigma | 10 μM | 30 min |
| **BMH-21** |  |  |  |  |
| **ML-60218** |  |  |  |  |

### Plasmids and transfections

For rDNA DSB induction by CRISPR/Cas9, four guide RNAs (gRNA) plasmids were used ^46^. Three of them were pMA plasmids expressing sequences targeting rDNA present on all five acrocentric short arms (Table 2) and the other was a control plasmid expressing the gRNA scaffold but no target sequence. Plasmids carry out ampicillin resistant marker.

**Table 2.**
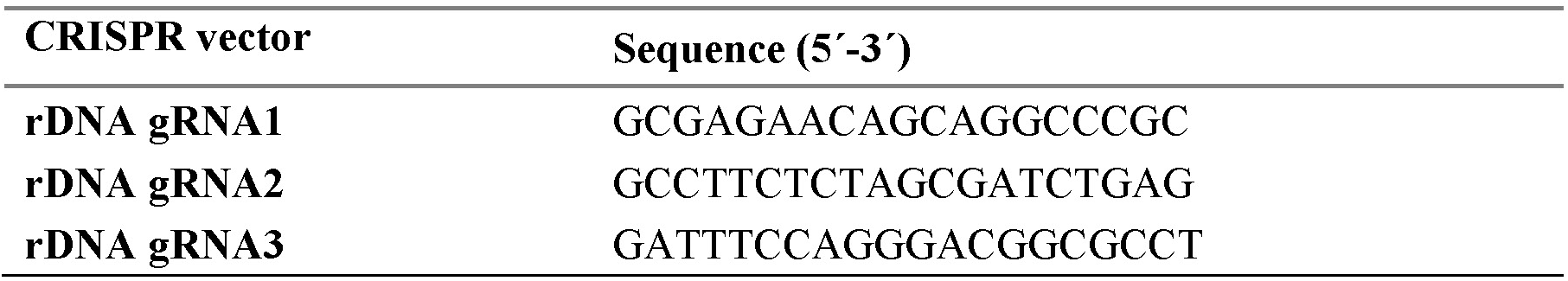
sgRNA sequences.

Transient transfection of U2OS-Cas9-NBS1-GFP cells was carried out using Lipofectamine LTX with plus reagent (Invitrogen) following manufacturer’s instructions. Briefly, cells were seeded in different plate formats, accordingly to the requirements of each specific experiment, to be 70-90% confluent at transfection. Then, the three gRNA plasmids pooled in a ratio 1:1:1 and PLUS reagent were diluted in Opti-MEM and mixed with Lipofectamine LTX Reagent blended into Opti-MEM, in a 1:1 ratio. Solution was incubated 5 min at room temperature and finally was added dropwise to the plate with gentle rocking. Cells were then collected 0-6 hours after transfection unless otherwise indicated.

### Inmunofluorescence

Cells were grown on sterile coverslips, transfected with gRNAs as previously described and treated or not with inhibitors as indicated in each figure. Cells were fixed with 4% paraformaldehyde (w/v) in PBS for 10 min, washed twice with PBS and then permeabilised with 0.2% Triton X-100 in PBS for 5 min. Following two washes with PBS, cells were blocked for at least 1 hour with 5% FBS diluted in PBS, incubated with adequate primary antibodies (Table 3) diluted in 5% FBS in PBS for 2 hours at room temperature, washed again with PBS and then, incubated with the appropriate secondary antibodies (Table 3) and 4′-6-diamidino-2-phenylindole (DAPI, 10 ng/ml) in blocking buffer for 1 hour at room temperature in the dark. Coverslips were mounted with Fluoromount-G mounting medium (Invitrogen), and images were acquired using a LEICA microscope. Image analysis was performed using ImageJ software by quantifying the number of nucleolar caps, nucleolar mean intensity, and fluorescence intensity profiles. Intensity profiles were obtained by drawing a line ROI across the image and using the ’Plot Profile’ tool to generate a graph of intensity values along the line. Experiments were repeated at least three times independently.

**Table 3:** Antibodies.

| Antibody | Species | Reference / Suppliers | Application |
| --- | --- | --- | --- |
| Nucleolin | Rabbit | Abcam; ab70493 | IF (1:1000) |
| Cas9 | Mouse | Abcam; ab191468 | WB (1:1000) |
| β-importin | Mouse | Abcam; ab2811 | WB (1:5000) |
| RNAPII | Mouse | Abcam; ab817 | IF (1:1000) |
| RNAPII S5P | Rabbit | Abcam; ab5131 | IF (1:2000) |
| RNAPII S2P | Rabbit | Abcam; ab5095 | IF (1:2000) |
| RNAPII pY1 | Mouse | Active Motif; 61383 | IF (1:1000) |
| BrdU | Rat | Bio-rad AbD Serotec; OBT0030 | IF (1:500) |
| RNAPIII | Mouse | Santa Cruz; SC-48365 | IF (1:500) |
| CtIP | Mouse | Active Motif; 61141 | IF (1:250)/WB (1:500) |
| 53BP1 | Rabbit | Novus, NB100-904 | IF (1:500) |
| BRCA1 | Mouse | Santa cruz; sc-6954 | IF 1:100 |
| H3K36me3 | Rabbit | Abcam ab9050 | IF 1:500 |
| HRP-conjugated goat anti-mouse IgG (H+L) | Goat | Jackson, 111-035-146 | WB (1:5000) |
| Alexa Fluor 647 anti-rabbit | Goat | Invitrogen, A21245 | IF (1:1000) |
| Alexa Fluor 546 anti-rabbit | Goat | Invitrogen, A11010 | IF (1:1000) |
| Alexa Fluor 546 anti-mouse | Goat | Invitrogen, A11003 | IF (1:1000) |

### High-content image acquisition

Quantitative image-based cytometry (QIBC) was performed as previously described ^12^. Briefly, the images were acquired in an automatic and unbiased way by scanR acquisition software 3.0 (Leica) and analysed by scanR image analysis software 3.0. The results were exported as txt files. The txt data set was further processed with spotfire and PRISM 8 software (Graphpad Software Inc) for further analysis. Statistical significance was determined with Ordinary one-way and two-way ANOVA tests using multiple comparisons using PRISM 8 software (Graphpad Software Inc).

### Metabolic labeling of nascent RNA by EU

Nascent RNA was visualized using metabolic labeling using Invitrogen™ Click-iT™ RNA Alexa Fluor™ 594 imaging kit, with modifications. Briefly, U2OS-Cas9-NBS1-GFP cells were seeded onto coverslips, transfected with gRNAs as previously described, treated 2 hours with 1 μM of THZ1 inhibitor 4 hours after transfection and pulsed for 30 min with EU at a final concentration of 1 mM. Following treatments, cells were fixed with 4% paraformaldehyde (PFA) for 10 min, permeabilised with 0.5% Triton X-100 and washed 3 times with Tris buffered saline (TBS) (50 mM Tris-HCl pH 8.0, 150 mM NaCl). Then, samples were incubated 30 min with fresh prepared Click- iT reaction cocktail, containing 5 µM Alexa Fluor 594 azide, 2 mM CuSO4 and 100 mM sodium ascorbate. After removal of the reaction cocktail, cells were washed three times with TBS and stained with DAPI. Finally, coverslips were mounted onto glass slides using Fluoromount-G (Invitrogen) and cells were imaged by fluorescence microscopy. Image analysis was performed by measuring the intensity profile using the ImageJ software drawing a line ROI over the image and using the "Plot Profile" tool to generate a graph of intensity values along that line.

### Immunoblotting

Whole-cell extracts were prepared in Laemmli buffer 2X (4% SDS, 20 % glycerol, 125 mM Tris−HCl, pH 6.8, 0.02% bromophenol blue, 200 mM DTT) by boiling at 100 °C for 10 min and sonicated in a Bioruptor for 2 cycles of 30 sec ON/30 sec OFF at high intensity. Proteins were resolved by SDS-PAGE and transferred to nitrocellulose membranes (AmershamTM Protran® 0.45 µm, GE Healthcare). Membranes were blocked for 1 hour with 5% BSA in TTBS (15 mM Tris- HCl, pH 7.5, 200 mM NaCl, 0.1% (v/v) Tween-20), followed by incubation with the appropriate primary antibodies overnight at 4°C (Table 3). After being washed with TTBS, membranes were incubated with HRP-conjugated secondary antibodies (Table 3) for 1 hour at room temperature. Chemiluminescence was performed using Clarity Western ECL Substrate (Bio-Rad), and results were acquired using the ChemiDoc Touch Imaging System (Bio-Rad) and visualised with Image Lab software 5.1 (Bio-Rad).

### Micronuclei assay

Cells were seeded onto coverslips, transfected with gRNAs as previously described and treated 2 hours with 1 μM of THZ1 inhibitor 4 hours after transfection. Following treatment, the medium was replaced by fresh one and cytochalasin B (Sigma-Aldrich) was added at 4 µg/ml. 20 hours post-treatment, cells were fixed in PBS-4% paraformaldehyde 10 min, permeabilised in PBS-0.2% Triton X-100 5 min, blocked 30 min in blocking buffer (PBS-5% FBS) and incubated 1-2 hours with nucleolin primary antibody (Table 3) in blocking buffer. Cells were then washed two times with PBS and incubated for 45 min with the corresponding secondary antibody (Table 3) diluted in blocking buffer plus DAPI (Sigma-Aldrich) in darkness. Finally, coverslips were washed twice with PBS, dried and mounted onto glass slides using Fluoromount-G (Invitrogen). Micronuclei containing or not nucleolin were counted only from binucleated cells.

### Clonogenic survival assay

For clonogenic survival assays, cells were transfected with gRNA as previously described for rDNA DSBs induction and 4 hours post-transfection cells were treated with different concentration (0.1 μM, 0.5 μM, 1μM) of THZ1 or vehicle (DMSO) as control for 2 hours. Then, cells were counted and re-seeded in 6-well plates at two different concentrations, 1000 and 500 cells per well, in triplicates. Cells were then grown at 37° C for 7-8 days to allow colonies formation. Afterwards, plates were washed in PBS and fixed and visualised by staining with 1% methylene blue diluted in 20% ethanol, followed by washes with water to remove the excess of staining. Plates were left to dry overnight and the number of colonies per well was scored for each condition and normalised to the untreated condition.

### MTT assay

MTT [3-(4,5-dimethylthiazol-2-yl)-2,5-diphenyltetrazolium bromide] assay (Roche) was performed following manufacturer’s indications. After transfection with gRNA and incubation with different concentration (0.1 μM, 0.5 μM, 1 μM) of THZ1 or DMSO for 2 hours as previously described, cells were seeded at a density of 2×10^3^ cells/well in 100 μl of medium on a 96-well plate in triplicates. After 5 days growing, 10 μl of MTT (5 mg/ml) was added to each well, and plates were incubated at 37°C for 4 hours. Then, 100 μl of 10% SDS was added, and cells were incubated overnight at 37°C to dissolve the insoluble purple formazan products. A570 was measured using a microplate reader with a reference wavelength of 690 nm.

### PLA assay

Proximity ligation assay (PLA) was performed using the Duolink In Situ Red Starter Kit Mouse/Rabbit (Sigma-Aldrich) according to the manufacturer’s instructions. First, U2OS-Cas9- NBS1-GFP cells were processed like immunofluorescence, blocked with blocking solution from the Duolink PLA Kit for 30 min at 37 °C and staining with specific primary antibodies (Table 3) overnight at 4° C. After that, cells were incubated with MINUS and PLUS secondary PLA probes for 1 h at 37 °C, and the detection of these probes was carried out using the ScanR microscope (Olympus). The number of PLA signals per nucleus was scored in at least 100 cells per sample.

## Acknowledgments

Thanks to Fernando Gómez Herrerośs lab for helpful discussions. Thanks to Diana Aguilar-Morante and Oliver Quevedo for technical advice and discussions.

## Funding

AECC-Investigator Grant (INVES20017GOME)

University of Seville-Plan Propio-Grant: 2021/00001263 (PV, GS) MSCA-IF-2017 (795930) PID2022-137280OB-I00 CNS2023-143939

R+D+i PID2022-136564NB-I00 (MCIN/AEI/10.13039/501100011033 and ERDF/UE "A way of making Europe")

Andalusian Regional Government (P20_00581, BIO-271) University of Seville (US-1380394, US/JUNTA/FEDER,UE). Novo Nordisk Foundation (NNF20OC0060590)

Danish Council for Independent Research (DFF-7016-00313) Swedish Research Council (VR-MH 2014-46602-117891-30) Danish National Research Foundation (project CARD, DNRF 125) Danish Cancer Society (R204-A12617-B153)

## Author contributions

Conceptualisation: DGC, JB Methodology: CCR, BSQ, OQ, LC, DGC Investigation: JCD, JB, DGC Supervision: DGC

Writing—original draft: DGC

Writing—review & editing: CCR, DHL, JCD, JB, DGC

## Data and materials availability

All data, code, and materials used in the analyses must be available in some form to any researcher for purposes of reproducing or extending the analyses.

## References

1. Zisi, A., Bartek, J. & Lindstrom, M.S. Targeting Ribosome Biogenesis in Cancer: Lessons Learned and Way Forward. Cancers (Basel) 14 (2022).

2. Derenzini, M. et al. Nucleolar function and size in cancer cells. Am J Pathol 152, 1291–1297 (1998).

3. Weeks, S.E., Metge, B.J. & Samant, R.S. The nucleolus: a central response hub for the stressors that drive cancer progression. Cell Mol Life Sci 76, 4511–4524 (2019).

4. Hilton, J. et al. Results of the phase I CCTG IND.231 trial of CX-5461 in patients with advanced solid tumors enriched for DNA-repair deficiencies. Nat Commun 13, 3607 (2022).

5. Lin, Y.L. & Pasero, P. Interference between DNA replication and transcription as a cause of genomic instability. Curr Genomics 13, 65–73 (2012).

6. Warner, J.R. The economics of ribosome biosynthesis in yeast. Trends Biochem Sci 24, 437–440 (1999).

7. Moss, T. & Stefanovsky, V.Y. At the center of eukaryotic life. Cell 109, 545–548 (2002).

8. Bowry, A., Kelly, R.D.W. & Petermann, E. Hypertranscription and replication stress in cancer. Trends Cancer 7, 863–877 (2021).

9. Kotsantis, P. et al. Increased global transcription activity as a mechanism of replication stress in cancer. Nat Commun 7, 13087 (2016).

10. Curti, L. et al. CDK12 controls transcription at damaged genes and prevents MYC-induced transcription-replication conflicts. Nat Commun 15, 7100 (2024).

11. Korsholm, L.M. et al. Recent advances in the nucleolar responses to DNA double-strand breaks. Nucleic Acids Res 48, 9449–9461 (2020).

12. Gomez-Cabello, D., Pappas, G., Aguilar-Morante, D., Dinant, C. & Bartek, J. CtIP-dependent nascent RNA expression flanking DNA breaks guides the choice of DNA repair pathway. Nat Commun 13, 5303 (2022).

13. Gomez-Cabello, D., Jimeno, S., Fernandez-Avila, M.J. & Huertas, P. New tools to study DNA double-strand break repair pathway choice. PLoS One 8, e77206 (2013).

14. Jackson, S.P. & Bartek, J. The DNA-damage response in human biology and disease. Nature 461, 1071–1078 (2009).

15. Heyer, W.D., Ehmsen, K.T. & Liu, J. Regulation of homologous recombination in eukaryotes. Annu Rev Genet 44, 113–139 (2010).

16. Lopez-Saavedra, A. et al. A genome-wide screening uncovers the role of CCAR2 as an antagonist of DNA end resection. Nat Commun 7, 12364 (2016).

17. Larsen, D.H. & Stucki, M. Nucleolar responses to DNA double-strand breaks. Nucleic Acids Res 44, 538–544 (2016).

18. van Sluis, M. & McStay, B. A localized nucleolar DNA damage response facilitates recruitment of the homology-directed repair machinery independent of cell cycle stage. Genes Dev 29, 1151–1163 (2015).

19. Kruhlak, M. et al. The ATM repair pathway inhibits RNA polymerase I transcription in response to chromosome breaks. Nature 447, 730–734 (2007).

20. Larsen, D.H. et al. The NBS1-Treacle complex controls ribosomal RNA transcription in response to DNA damage. Nat Cell Biol 16, 792–803 (2014).

21. Lesage, E., Clouaire, T. & Legube, G. Repair of DNA double-strand breaks in RNAPI- and RNAPII-transcribed loci. DNA Repair (Amst) 104, 103139 (2021).

22. Gal, Z. et al. Hyper-recombination in ribosomal DNA is driven by long-range resection-independent RAD51 accumulation. Nat Commun 15, 7797 (2024).

23. Liu, S. et al. RNA polymerase III is required for the repair of DNA double-strand breaks by homologous recombination. Cell 184, 1314–1329 e1310 (2021).

24. Domingo-Prim, J. et al. EXOSC10 is required for RPA assembly and controlled DNA end resection at DNA double-strand breaks. Nat Commun 10, 2135 (2019).

25. Burger, K., Schlackow, M. & Gullerova, M. Tyrosine kinase c-Abl couples RNA polymerase II transcription to DNA double-strand breaks. Nucleic Acids Res 47, 3467–3484 (2019).

26. Vitor, A.C. et al. Single-molecule imaging of transcription at damaged chromatin. Sci Adv 5, eaau1249 (2019).

27. Pappas, G. et al. MDC1 maintains active elongation complexes of RNA polymerase II. Cell Rep 42, 111979 (2023).

28. Cohen, S. et al. Senataxin resolves RNA:DNA hybrids forming at DNA double-strand breaks to prevent translocations. Nat Commun 9, 533 (2018).

29. Ouyang, J. et al. RNA transcripts stimulate homologous recombination by forming DR- loops. Nature 594, 283–288 (2021).

30. Bonath, F., Domingo-Prim, J., Tarbier, M., Friedlander, M.R. & Visa, N. Next-generation sequencing reveals two populations of damage-induced small RNAs at endogenous DNA double-strand breaks. Nucleic Acids Res 46, 11869–11882 (2018).

31. Abraham, K.J. et al. Nucleolar RNA polymerase II drives ribosome biogenesis. Nature 585, 298–302 (2020).

32. Khosraviani, N. et al. Nucleolar Pol II interactome reveals TBPL1, PAF1, and Pol I at intergenic rDNA drive rRNA biogenesis. Nat Commun 15, 9603 (2024).

33. Trifault, B. et al. Nucleolar detention of NONO shields DNA double-strand breaks from aberrant transcripts. Nucleic Acids Res 52, 3050–3068 (2024).

34. Korsholm, L.M. et al. Double-strand breaks in ribosomal RNA genes activate a distinct signaling and chromatin response to facilitate nucleolar restructuring and repair. Nucleic Acids Res 47, 8019–8035 (2019).

35. Nilson, K.A. et al. THZ1 Reveals Roles for Cdk7 in Co-transcriptional Capping and Pausing. Mol Cell 59, 576–587 (2015).

36. Olson, C.M. et al. Pharmacological perturbation of CDK9 using selective CDK9 inhibition or degradation. Nat Chem Biol 14, 163–170 (2018).

37. Aoi, Y. et al. SPT5 stabilization of promoter-proximal RNA polymerase II. Mol Cell 81, 4413–4424 e4415 (2021).

38. Wang, Y., Lu, J.J., He, L. & Yu, Q. Triptolide (TPL) inhibits global transcription by inducing proteasome-dependent degradation of RNA polymerase II (Pol II). PLoS One 6, e23993 (2011).

39. Meisenberg, C. et al. Repression of Transcription at DNA Breaks Requires Cohesin throughout Interphase and Prevents Genome Instability. Mol Cell 73, 212–223 e217 (2019).

40. Carvalho, S. et al. SETD2 is required for DNA double-strand break repair and activation of the p53-mediated checkpoint. Elife 3, e02482 (2014).

41. Markert, J.W., Soffers, J.H. & Farnung, L. Structural basis of H3K36 trimethylation by SETD2 during chromatin transcription. Science 387, 528–533 (2025).

42. Warmerdam, D.O., van den Berg, J. & Medema, R.H. Breaks in the 45S rDNA Lead to Recombination-Mediated Loss of Repeats. Cell Rep 14, 2519–2527 (2016).

43. Wang, K. et al. Blockage of Autophagic Flux and Induction of Mitochondria Fragmentation by Paroxetine Hydrochloride in Lung Cancer Cells Promotes Apoptosis via the ROS- MAPK Pathway. Front Cell Dev Biol 7, 397 (2019).

44. Hatch, E.M., Fischer, A.H., Deerinck, T.J. & Hetzer, M.W. Catastrophic nuclear envelope collapse in cancer cell micronuclei. Cell 154, 47–60 (2013).

45. Saur, F. et al. Transcriptional repression facilitates RNA:DNA hybrid accumulation at DNA double-strand breaks. Nat Cell Biol (2025).

46. van Sluis, M. & McStay, B. Nucleolar reorganization in response to rDNA damage. Curr Opin Cell Biol 46, 81–86 (2017).

47. Imrichova, T. et al. Dynamic PML protein nucleolar associations with persistent DNA damage lesions in response to nucleolar stress and senescence-inducing stimuli. Aging (Albany NY) 11, 7206–7235 (2019).

48. Hornofova, T. et al. Phospho-SIM and exon8b of PML protein regulate formation of doxorubicin-induced rDNA-PML compartment. DNA Repair (Amst) 114, 103319 (2022).

49. Urbancokova, A. et al. Topological stress triggers persistent DNA lesions in ribosomal DNA with ensuing formation of PML-nucleolar compartment. Elife 12 (2024).

