## Supplementary material for "RNA Polymerase II-mediated transcription is required for repair of ribosomal DNA breaks in nucleolar caps, guarding against genomic instability": Figure Legends

**Fig. 1. RNAPII, but not RNAPI or RNAPIII, inhibition significantly impairs nucleolar cap formation. (a)** Schematic representation of CRISPR/Cas9-induced DSBs in rDNA. The three designed gRNAs specifically target the 5′ ETS or the central region of the IGS (highlighted in red). **(b)** The percentage of cells with more than one nucleolar caps upon DSBs induction and RNA polymerases inhibition is plotted. Cells were transfected with gRNA targeting rDNA, and 3–4 h later were exposed to BMH 21 (RNAPI inhibitor, 1 μM, 3 h), THZ1 (RNAPII inhibitor, 1 μM, 2 h), ML 60218 (RNAPIII inhibitor, 100 μM, 3 h), or DMSO (vehicle control). Graph represents the average and SD of six independent experiments. **(c)** Number of nucleolar caps per cell in the same conditions as in (b). **(d)** Scatter plot showing the number of nucleolar caps per cell under the same conditions as in (c), from one representative experiment. **(e)** Representative fluorescence images of NBS1-GFP (green) and RNAPI (red) in U2OS-NBS1-GFP-Cas9 cells treated under the same conditions as in (b), along with a representative fluorescence intensity profile across the indicated line. Colocalization was assessed by measuring fluorescence intensity along the line. Nuclei were stained with DAPI**. (f)** Percentage of nucleolar caps showing colocalization of NBS1-GFP and RNAPI, measured as shown in (e). At least 60 nucleolar caps from three biologically independent experiments were analyzed. **(g)** Same as (e) but for RNAPII phosphorylated in Serine-5 (RNAPII-S5P; red). **(h)** Same as (f) but for NBS1-GFP and RNAPII-S5P colocalization.
Statistical significance was calculated with one-way anova.

**Fig. 2. RNAPII transcription initiation and elongation are essential to accurate rDNA repair. (a)** Schematic illustration of the experimental timeline. U2OS-NBS1-GFP-Cas9 cells were seeded at the start of experiments. The following day, cells were transfected with gRNA targeting rDNA or the control empty vector for 6 hours (green box). 4 hours post-transfection cells were treated with THZ1 for 2 or 24 hours (orange box). In the case of 24h-treatment, media was replaced after 2 hours to eliminate the transfection agents and THZ1 was added again in the fresh medium. Finally, cells were collected and differently processed depending on the experiment. **(b)** Representative fluorescence microscopy images of RNAPII phosphorylated in Serine-5 (RNAPII-S5P; red) after gRNA transfection and THZ1 treatment. Nuclei were stained with DAPI. **(c)** Quantification of RNAPII-S5P intensity in cells from (b). A representative experiment from 3 replica is shown. At least 1000 cells were quantified. **(d)** Quantification of BrdU staining under non-denaturing conditions to mark ssDNA as a DNA resection marker in the same conditions as in (b). **(e)** Representative microscopy images of nucleolar caps marked with NBS1-GFP in U2OS-NBS1-GFP-Cas9 following rDNA DSBs induction by gRNA transfection

and RNAPII inhibition by THZ1 treatment. Nuclei were stained with DAPI and nucleoli with anti-nucleolin antibody. **(f)** The percentage of cells with more than two nucleolar caps upon DSBs induction and RNAPII inhibition is plotted. Graph represents the average and SEM of 3 independent experiments. At least 1000 cells were count. **(g)** Number of nucleolar caps per cell in the same conditions as in (f). A representative experiment of 3 is shown. **(h)** Cas9 levels analysed by western blotting from whole protein extract of U2OS-NBS1-GFP-Cas9 cells transfected with gRNA and treated with THZ1. β-importin was used as a loading control. **(i)** Representative image of RNAPII transcription cycle for showing assembly, initiation and pause, re-initiation and elongation-termination stages. Phosphorylation of different residues are marked: Serine-5 phosphorylation (red circles), Serine-2 phosphorylation (yellow circles) and Tyrosine 1 phosphorylation (orange circles). Different RNAPII inhibitors with their targets are shown. **(j)** Representative fluorescence images of NBS1-GFP nucleolar caps (green) and RNAPII S5P (red) of U2OS-NBS1-GFP-Cas9 cells transfected with the gRNA or the empty control and treated with the different RNAPII inhibitors indicated.
**(k)** Percentage of cells in the conditions of (j) with more than two nucleolar caps. Graph represents the average and SEM of 3 independent experiments. At least 1000 cells were count. **(l)** Number of nucleolar caps per cell in the same conditions as in (j). A representative experiment of 3 replicas is shown. **(m)** Graph shows nuclear quantification of RNAPII S5P intensity under the experimental conditions cued in (j). At least 1000 cells were quantified. The average and SEM of 3 experiments is shown.
Statistical significance was calculated with Ordinary One-way Anova.

**Fig. 3. RNAPII colocalises with nucleolar caps. (a)** Representative fluorescence images for NBS1-GFP (green) and RNAPII-S5P (red) fluorescence from U2OS-NBS1-GFP-Cas9 cells transfected with gRNA to induce rDNA DSBs. Right zoomed-in images show in detail the position of line scanned for fluorescent intensity measurement. Representative intensity profile for NBS1-GFP and RNAPII-S5P fluorescence across the line is shown. At least 10 images of 3 biologically independent experiments were quantified. **(b)** Same as (a) but for RNAPII-S2P (red). **(c)** Same as (a) but for RNAPII-Y1P (red).

**Fig. 4. DNA resection is affected by RNAPII inhibitor in the nucleolar cap. (a)** Representative fluorescence images of NBS1-GFP (green) and CtIP (red) in U2OS-NBS1-GFP-Cas9 cells transfected with gRNA to induce rDNA DSBs and exposed to different RNAPII inhibitors (THZ1 1 μM, 2 h; DRB 100 μM, 2 h; Triptolide 10 μM, 30 min) or DMSO (vehicle control). Representative intensity profile for NBS1-GFP and CtIP fluorescence across the line is shown. Nuclei were stained with DAPI. **(b)** Percentage of nucleolar caps showing colocalization of NBS1-GFP and CtIP, measured as shown in (a). At least 60 nucleolar caps from three biologically independent experiments were analyzed. Statistical significance was calculated with Ordinary One-way Anova. **(c)** Graph shows the average number of NBS1-CtIP PLA foci (Proximity ligation assay) for protein interactions among NBS1 and CtIP after gRNA transfection and THZ1 treatment. At least 1000 cells of 3 independent experiments were analysed. Statistical significance was calculated with unpaired t test. **(d)** Representative fluorescence images of NBS1-GFP (green) and BrdU (red) colocalisation upon gRNA transfection, THZ1 treatment and under non-denaturing conditions. Representative fluorescence intensity profiles for NBS1-GFP (green) and BrdU (red) quantified along the traced line is shown. Nuclei were stained with DAPI. At least 10 images of 3 biologically independent experiments were quantified. **(e)** Same as (d) but for EU (red).

**Fig. 5. Ribosomal Biogenesis is not affected by THZ1 treatment at 6 and 24 hours. (a)** Polysome profiles on sucrose gradients of U2OS-NBS1-GFP-Cas9 cells upon gRNA transfection and THZ1 treatment for 2 hours (left panels) or 24 hours (right panels). The peaks of free 40S and 60S r-subunits, monosomes 80S and polysomes are indicated. **(b)** Schematic representations of pre-rRNA intermediates in human cells, with the processing/cleavage sites, ETS and ITS, indicated. The positions of the probes used for northern blot hybridizations are marked with colored lines and letters (A-H). Modified from (2). **(c)** Representative image of accumulation levels of rRNA precursors in U2OS-NBS1-GFP-Cas9 cells upon gRNA transfection and THZ1 treatment. Total RNA extracted from each sample was analyzed by northern blotting. The RNA molecules detected using radiolabeled oligonucleotide probes are indicated on the right. **(d)** Quantification of rRNA precursors accumulation levels in the same conditions as (c). The mean and SEM of three different experiment is shown in each graph.

**Fig. 6. RNAPII and BRCA1 recruitment at Nucleolar caps is regulated by CtIP. (a)** Representative fluorescence images showing colocalization of NBS1-GFP (green) and RNAPII (red) upon gRNA transfection and CtIP knockdown, along with fluorescence intensity profiles quantified along the indicated line. Right zoomed-in images show the region where fluorescence intensity was measured along the drawn line. At least 30 images from 3 biologically independent experiments were analysed. **(b)** Percentage of NBS1-GFP and RNAPII colocalization upon gRNA transfection and CtIP knockdown. Data represent analysis of at least 60 nucleolar caps per sample. **(c)** Representative

Western blot showing CtIP knockdown efficiency after siRNA transfection. Tubulin was used as a loading control. **(d)** Same at (a) but for BRCA1 (red). **(e)** Same as (b) for NBS1-GFP and BRCA1 colocalization. **(f)** Same as (a) but for 53BP1 (red). **(g)** Same as (b) for NBS1-GFP and 53BP1 colocalization. Statistical significance was calculated with unpaired t test.

**Fig. 7. CtIP reduces H3K36me3 at nucleolar caps after rDNA damage, influencing repair pathway choice. (a)** Representative fluorescence images showing colocalization of NBS1-GFP (green), RNAPII (red) and H3K36me3 (gray) upon gRNA transfection and CtIP knockdown, along with fluorescence intensity profiles quantified along the indicated line. Right zoomed-in images show the region where fluorescence intensity was measured along the drawn line. At least 30 images from 3 biologically independent experiments were analysed. **(b)** Percentage of NBS1-GFP and H3K36me3 colocalization upon gRNA transfection and CtIP knockdown. Data represent analysis of at least 60 nucleolar caps per sample. **(c)** Same as (b) for NSB1-GFP, H3K36me3 and RNAPII colocalization. **(d)** Same as (a) but for BRCA1 (red). **(e)** Same as (c) for NBS1-GFP, H3K36me3 and BRCA1 colocalization. **(f)** Model in which CtIP enable RNAPII access to damaged rDNA regions, promoting H3K36me3 deposition and BRCA1 loading within nucleolar caps. Statistical significance was calculated with unpaired t test.

**Fig. 8. RNAPII inhibition affects cell viability and rDNA stability. (a)** Percentage of clonogenic cell survival of U2OS-NBS1-GFP-Cas9 cells following rDNA DSBs induction and THZ1 treatment compared with the undamaged and untreated control (Empty – DMSO). Cells were transfected with a vector expressing a guide RNA targeting the rDNA (gRNA) or an empty vector as a control (Empty) and treated with different concentrations (0.1 μM, 0.5 μM, 1 μM) of THZ1 or DMSO as control for 2 hours. 7-8 days later, clones were stained with methylene blue. In each experiment, the average number of colonies per condition was scored in triplicates. The average and SEM of four independent experiments are plotted. **(b)** Cell viability measured by MTT assay upon transfection and THZ1 treatment as in (a) and after 5 days of growing following MTT treatment. Graph represents the mean and SEM of 3 independent experiments performed with technical triplicates, relative to undamaged and untreated control. **(c)** Percentage of cells with micronuclei containing or not nucleolin, after gRNA transfection and 2 hours of 1 μM THZ1 treatment. Micronuclei was scored in 500 binucleated U2OS-NBS1-GFP-Cas9 cells per experiment. The average and SEM of 3 independent experiments is shown. **(d)** Representative fluorescence microscopy images of U2OS-NBS1-GFP-Cas9 cells with micronuclei after DSBs induction and THZ1 treatment. DNA is shown in blue, nucleolin in red and NBS1-GFP in green. An example of micronucleus is indicated by a full white arrow. Statistical significance was calculated with two-way ANOVA

---

**Fig. S1. Repair-associated nucleolar caps differ from RNAPI inhibition-induced caps. (a)** Representative fluorescence microscopy images of NBS1-GFP (green) and UBF (red) in U2OS-NBS1-GFP-Cas9 cells treated with RNAPI inhibitors (ActD, BMH-21, or CX5461), RNAPII inhibitor (THZ1), DMSO (vehicle control), or transfected with a gRNA targeting rDNA to induce rDNA DSBs or an empty vector. Nuclei were stained with DAPI and nucleoli with nucleolin. **(b)** Percentage of cells with more than one UBF-positive nucleolar cap under the same conditions as in (a). **(c)** Number of UBF-positive nucleolar caps per cell in the same conditions as in (a). **(d)** Percentage of cells with more than one NBS1-positive nucleolar cap under the same conditions as in (a). **(e)** Number of NBS1-positive nucleolar caps per cell in the same conditions as in (a).

**Fig. S2. Partial colocalization of RNAPIII in nucleolar caps. (a)** Representative fluorescence images of NBS1-GFP (green) and RNAPIII (red) in U2OS-NBS1-GFP-Cas9 cells transfected with gRNA targeting rDNA, and 3–4 h later were exposed to BMH 21 (RNAPI inhibitor, 1 µM, 3 h), THZ1 (RNAPII inhibitor, 1 µM, 2 h), ML-60218 (RNAPIII inhibitor, 100 µM, 3 h), or DMSO (vehicle control). Representative intensity profile for NBS1-GFP and RNAPIII fluorescence across the line is shown. Nuclei were stained with DAPI**. (b)** Percentage of nucleolar caps showing colocalization of NBS1-GFP and RNAPIII, measured as shown in (a). At least 60 nucleolar caps from three biologically independent experiments were analyzed. **(c)** Representative fluorescence images for NBS1-GFP (green) and RNAPIII (red). Zoomed-in images show in detail the position of line scanned for fluorescent intensity measurement, and representative intensity profile is shown.

**Fig. S3. THZ1 time-course and triptolide treatment effects on nucleolar cap formation. (a)** Percentage of U2OS-NBS1-GFP-Cas9 cells with more than two nucleolar caps at different time points after transient gRNA transfection, direct transfer of ectopic recombinant Cas9 protein and THZ1 treatment immediately after transfection. Graph represent the mean and SEM of three different experiment. **(b)** Nucleoplasm quantification of NBS1-GFP intensity under the conditions described in (a). At least 1000 cells were analysed. **(c)** Same as (b) but for RNAPII S5P marker. **(d)** Percentage of

U2OS-NBS1-GFP-Cas9 cells with more than one nucleolar cap 6 hours post-transfection with gRNA or the empty control and treated 30 minutes with 10 µM of RNAPII inhibitor Triptolide. Graph represents the average and SEM of 3 independent experiments. At least 500 cells were count. **(e)** Number of nucleolar caps per cell in the same conditions as in (d). A representative experiment of 3 is shown. Statistical significance was calculated with Ordinary One-way Anova (* $p<0.05$; ** $p<0.01$; *** $p<0.001$; **** $p<0.0001$).

**Fig. S4. Effects of THZ1 treatment on DSB repair proteins. (a)** CtIP and Cas9 protein levels analysed by western blot from whole protein extract of U2OS-NBS1-GFP-Cas9 cells transfected with gRNA and treated 30 minutes or 2 hours with THZ1. β-importin was used as a loading control. **(b)** Representative fluorescence images of NBS1-GFP (green) and 53BP1 (red) in U2OS-NBS1-GFP-Cas9 cells transfected with gRNA to induce rDNA DSBs and treated with THZ1 2 h. Representative intensity profile for NBS1-GFP and 53BP1 fluorescence across the line is shown. Nuclei were stained with DAPI and nucleolus with anti-nucleolin antibody. **(c)** Percentage of nucleolar caps showing colocalization of NBS1-GFP and 53BP1, measured as shown in (b). At least 60 nucleolar caps from three biologically independent experiments were analyzed. Statistical significance was calculated with Ordinary One-way Anova.

**Fig. S5. rDNA damage increases the levels of H3K36me3 nucleolar. (a)** Representative fluorescence images showing NBS1-GFP (green) and H3K36me3 (red) intensity in nucleolus selected by nucleophosmin (gray) upon gRNA transfection. At least 30 images from 3 biologically independent experiments were analysed. **(b)** Mean intensity of H3K36me3 in nucleolus upon gRNA transfection. Data represent of at least 500 cells per sample. Statistical significance was calculated with unpaired t test.

**Fig. S6. THZ1 treatment provokes rDNA instability after rDSBs.**
**(a)** Percentage of clonogenic cell survival of U2OS-NBS1-GFP-Cas9 cells after gRNA or the empty vector transfection and treatment with different concentrations of THZ1 or DMSO as control, relative to each untreated control (Empty – DMSO or gRNA – DMSO). The average and SEM of four independent experiments are plotted. In each experiment, the average number of colonies per condition was scored in triplicates. **(b)** MTT assay upon gRNA transfection and THZ1 long treatment at different concentrations. MTT treatment was performed after 5 days of growing. THZ1 was added 4 hours post-transfection and it was maintained during the 5 days of growth. Graph represents the mean and SEM of three independent experiments performed with technical triplicates,

relative to undamaged and untreated control. **(c)** Cells transfected and treated with different concentrations of THZ1 for 2 hours were stained with MitoTracker Deep Red dye. Flow cytometry-based analysis of MitoTracker fluorescence intensity relative to undamaged and untreated cells is shown. Values represent mean and SEM of four independent experiments. **(d)** Representative flow cytometry plots showing mitotracker intensity in the same conditions as (c). **(e)** Same as (c) but after 24 hours of THZ1 treatment. **(f)** Percentage of cells with micronuclei containing or not nucleolin, after gRNA transfection and 24 hours of 1 uM THZ1 treatment. Micronuclei was scored in 500 binucleated cells per experiment. The average and SEM of three independent experiments is shown. Statistical significance was calculated with two-way ANOVA (* $p<0.05$; ** $p<0.01$; *** $p<0.001$).
