## Supplementary material for "RNA Polymerase II-mediated transcription is required for repair of ribosomal DNA breaks in nucleolar caps, guarding against genomic instability": Suppl. Figures

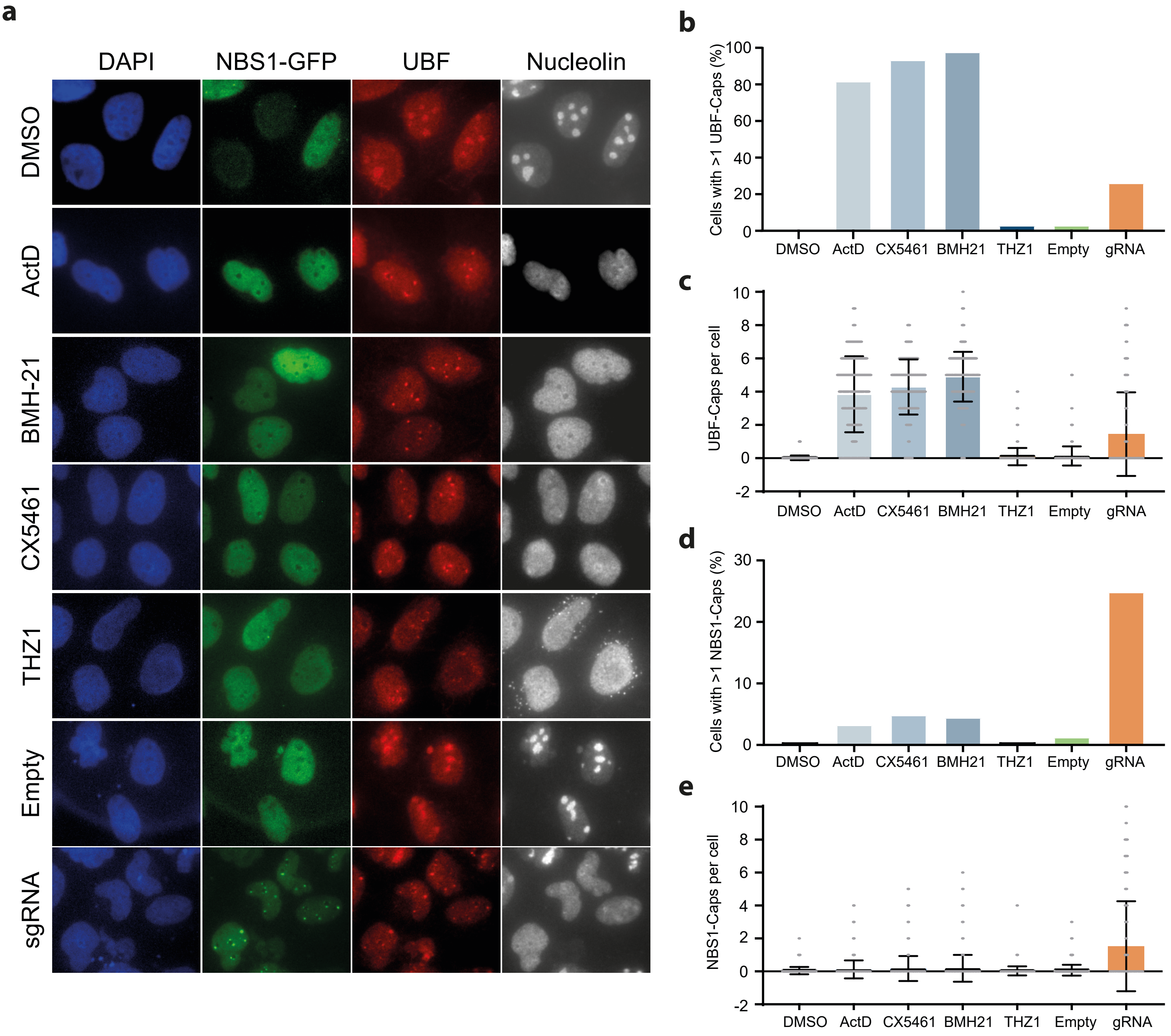


**Fig. S1.** Checa-Rodríguez, et al.

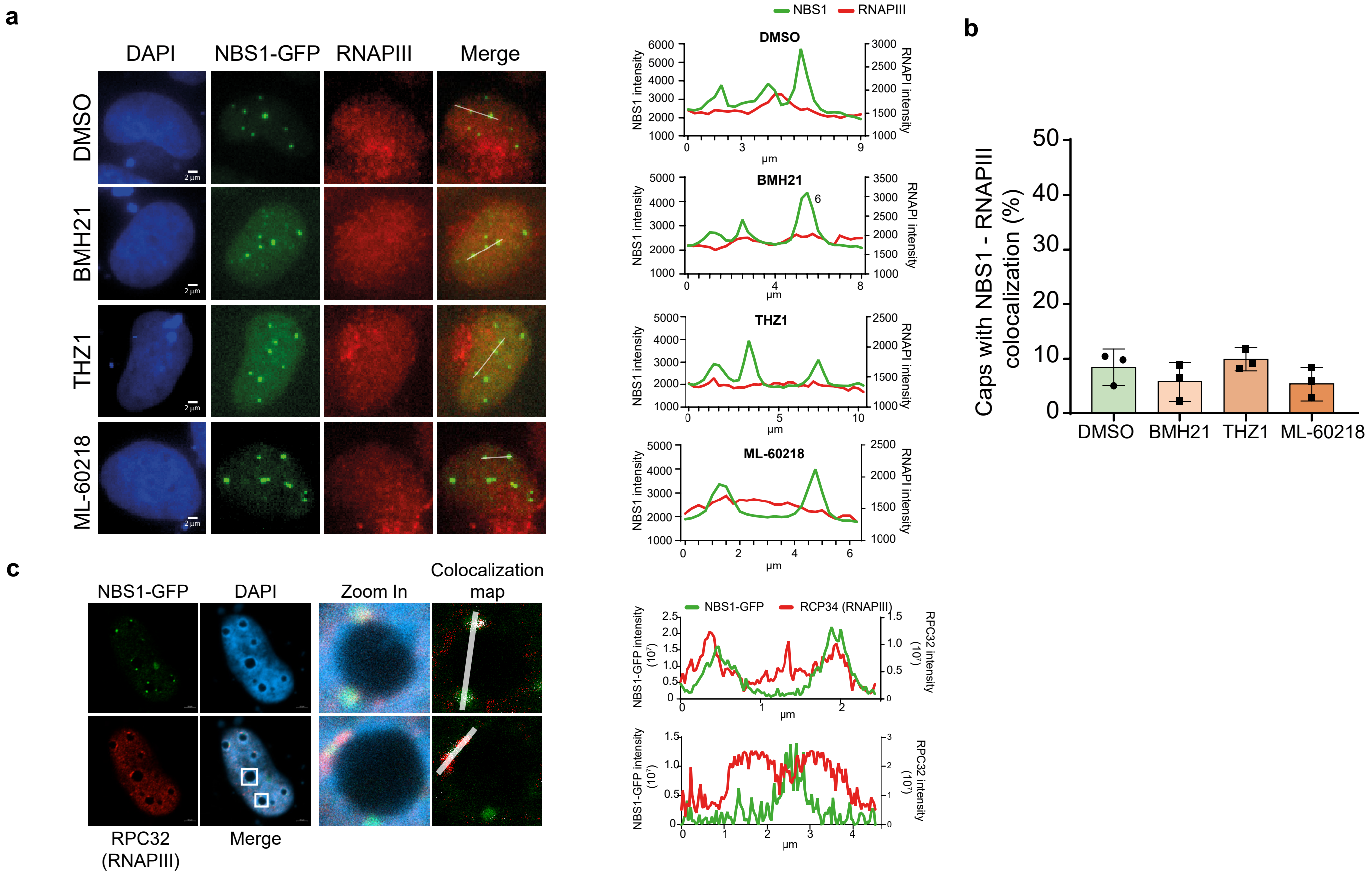


**Fig. S2.** Checa-Rodríguez, et al.

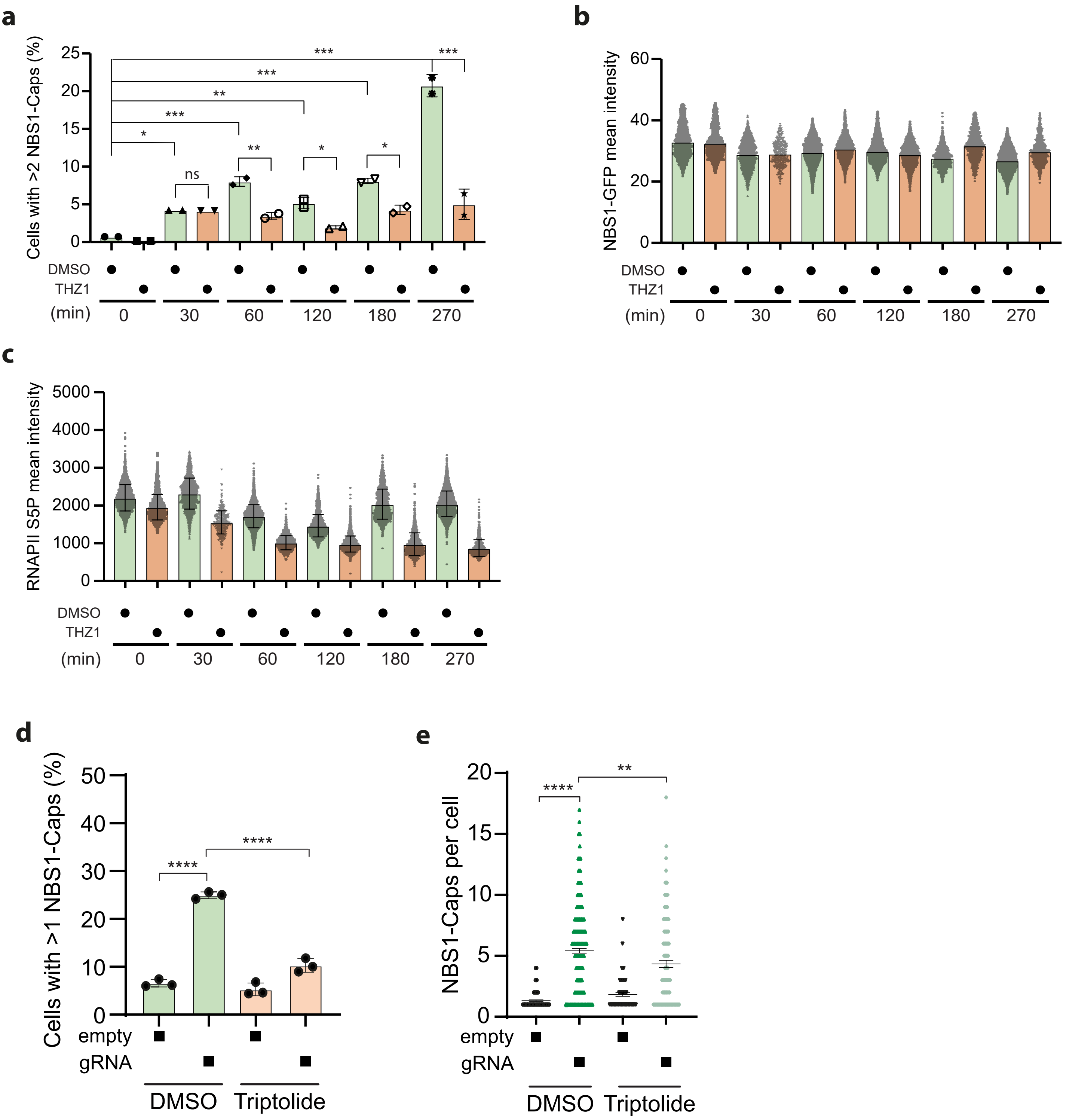


**Fig. S3.** Checa-Rodríguez, et al.

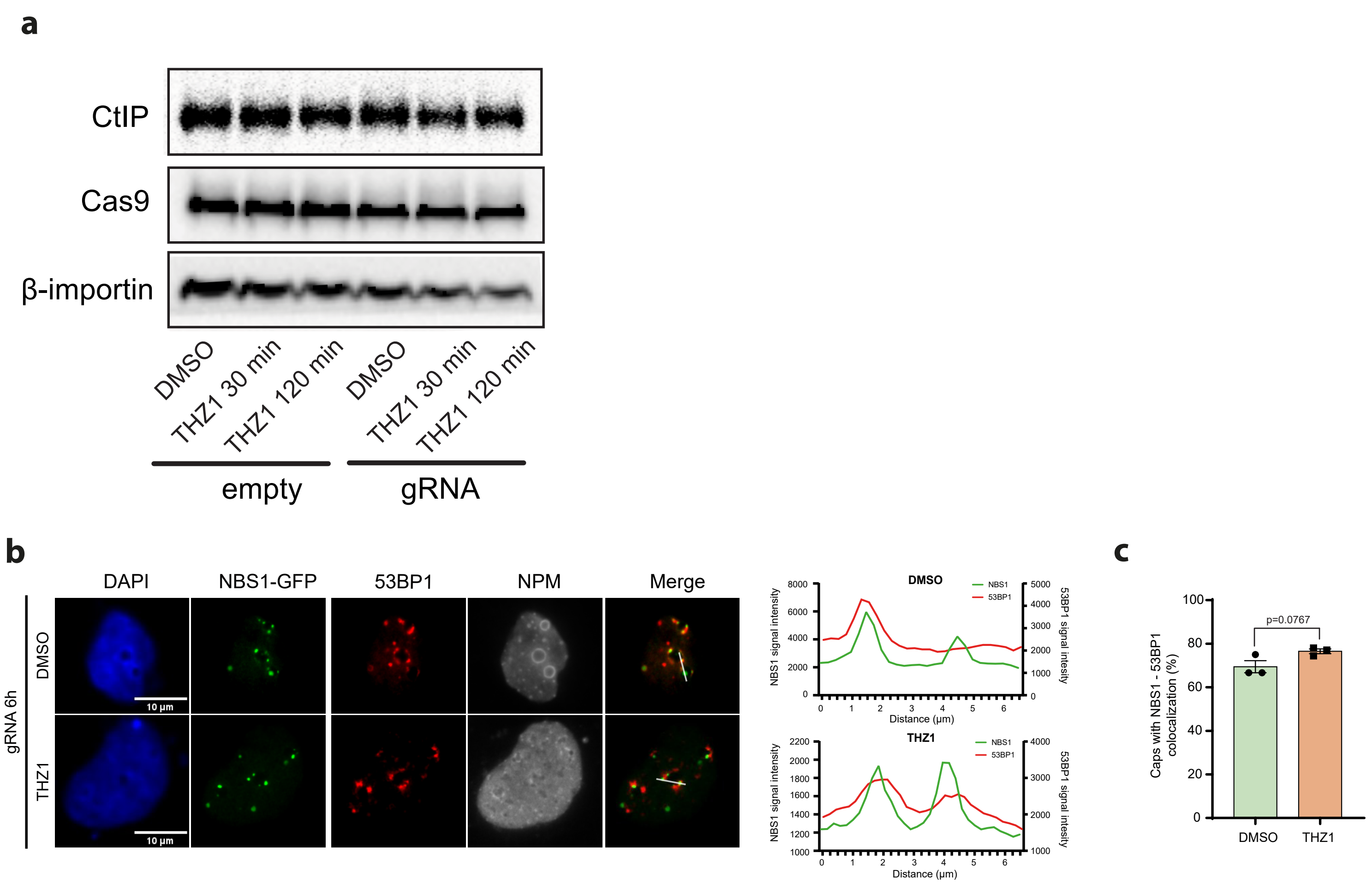


**Fig. S4.** Checa-Rodríguez, et al.

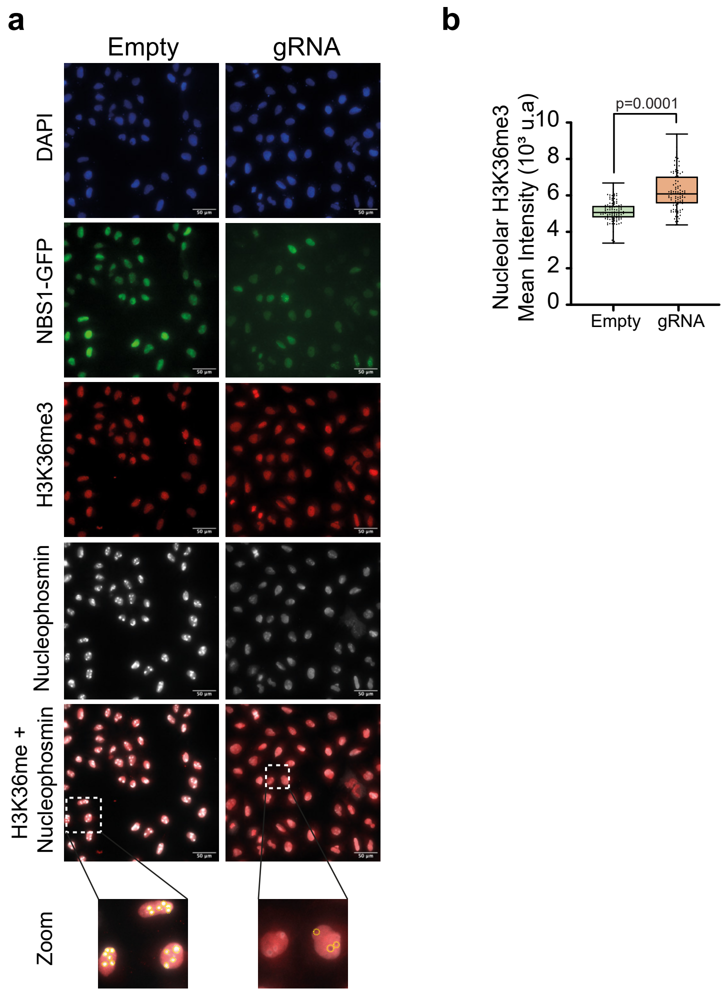


**Fig. S5.** Checa-Rodríguez, et al.

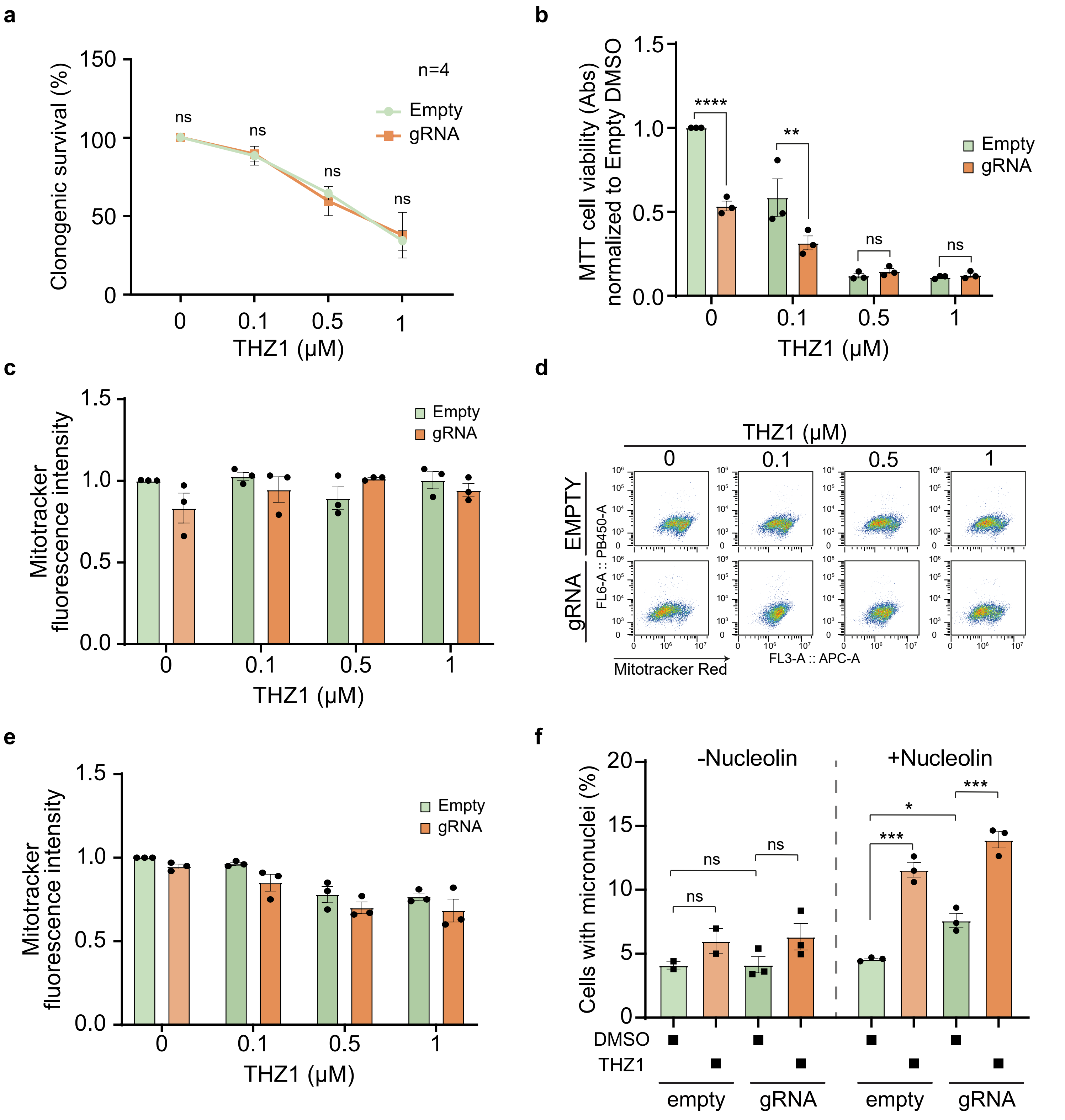


**Fig. S6.** Checa-Rodríguez, et al.
